# Ventromedial striatal GABAergic interneurons sex-dependently gate cost-benefit choices between food and exercise

**DOI:** 10.64898/2026.03.22.713369

**Authors:** Imane Hurel, Rim Fayad, Bastien Redon, Doriane Gisquet, Francisca Julio-Kalajzic, Abel Eraso-Pichot, Thierry Lesté-Lasserre, Astrid Cannich, Luigi Bellocchio, Giovanni Marsicano, Francis Chaouloff

**Author notes:** Life and Health Sciences Research Institute (ICVS), School of Medicine, University of Minho, Braga, Portugal. Joint last authors (Giovanni Marsicano, Francis Chaouloff).

## Abstract

Healthy energy balance relies on an equilibrated trade-off between the respective drives for food and exercise. However, the motivation circuitry underlying the choice between these two rewards remains unknown. Here, we developed a neuroeconomic model wherein mice living in operant chambers needed to choose between standard food and wheel running under increasing effort demands. Reward seeking resistance to increasing costs was then quantified using feeding and exercise essential values (EVs). Through conditional genetics and viral approaches, we show that cannabinoid type-1 receptors (CB_1_Rs) located on GABAergic neurons gate in necessary and sufficient manners exercise EV, exercise preference over feeding and exercise duration per rewarded sequence. Further, we report that these GABAergic neurons are located in the ventromedial striatum in males, but not females. CB_1_R deletion from medium spiny neurons did not impact exercise motivation, revealing an unforeseen role for ventromedial striatal GABAergic interneurons in effort-based decision-making between food and exercise.

## INTRODUCTION

Chronic eating disorders are most often depicted as dramatic excesses (e.g. obesity) or reductions (e.g. restricted type of anorexia nervosa) in food consumption. This shortcut however omits that these consumption alterations are first accounted for by imbalances in feeding motivation. This distinction is essential because the motivation and consumption dimensions of reward processes lie on distinct, albeit complementary, brain circuitries (*1*, *2*). Such a distinction extends to another rewarding activity, namely exercise (*3*), the motivation for which is most often oppositely altered in the two feeding pathologies mentioned above (*4–6*).

As opposed to the neurobiological substrates of eating motivation (*7*, *8*), those on which relies the motivation for exercise have been scarcely investigated although exercise (e.g. wheel running) is a strong reinforcer in laboratory rodents (*9–12*). As an illustration, free, and to a higher extent, conditioned access to running wheels, triggers synaptic changes (*13*) reminiscent of those measured with other natural and drug reinforcers (*14*). One limit in our knowledge of the bases of exercise motivation lies in the general use of free wheel running performance as a proxy for motivation (*15*). However, as for other rewards (*1*, *2*), the measure of exercise consumption (i.e. performance), fails to provide an adequate index of exercise motivation. Using instrumental conditioning wherein wheel running access obeyed nose poking under fixed ratio (FR) and progressive ratio (PR) reinforcement schedules, we have shown that cannabinoid type-1 receptor (CB_1_R)-expressing GABAergic neurons (likely located in the ventrotegmental area; VTA) control exercise motivation (*12*). Of note, this control did not extend either to the running performance per rewarded sequence (confirming the abovementioned dichotomy between the CNS bases of reward motivation and consumption) or to the motivation for another reward, namely palatable food. At first glance, these findings might be considered a breakthrough in the identification of the neurobiological bases of the balance between food and exercise drives. However, like most instrumental studies, animals underwent daily short experimental sessions before returning to their home cages under free feeding conditions. Besides questioning their relevance to chronic human situations (*16*), these temporary “closed economy” sessions (i.e. the sole daily periods during which mice were confronted to a running wheel and palatable food) differ from the human closed economy condition where effort- and value-based decision making for reward access is permanent. The use of similar reinforcer magnitudes for the comparison of rewards with different incentive valences is another limit. To overcome these translational weaknesses, we developed a unique closed economy paradigm in which mice live for days in operant chambers with food and exercise accesses governed by FR schedules. To further gain insight into the mechanisms gating the respective demand elasticities for food and exercise, the FR reinforcement schedules were gradually increased. These settings allowed to calculate for each reward its essential value (EV), an index of the effort-driven inelasticity of the reward demand (*17–19*). Of major importance, the EV for a reward is independent from its reinforcer magnitude (*18*, *20*, *21*), hence allowing a direct comparison between the EVs of different rewards (*20*). This so-called “neuroeconomic” paradigm thus permits an unbiased investigation on the motivation-based preference for one reward over another one and thereby its neurobiological bases.

This study used conditional gene deletions/re-expressions and viral approaches in male and female mice to reveal that CB_1_Rs, by virtue of 2-arachidonoylglycerol (2-AG) signaling, are essential for running, but not feeding, motivation. We then show that these CB_1_Rs are expressed by GABAergic neurons and that they are both necessary and sufficient for the control of exercise EV and exercise preference over food in both sexes. However, their brain region location is sex-specific as CB_1_Rs expressed by ventromedial striatal GABAergic interneurons control exercise motivation in males, but not females.

## RESULTS

### Development of a paradigm to measure the respective demand elasticities for food and exercise

To validate our permanent neuroeconomic paradigm (Fig. 1A), C57Bl/6N mice of both sexes were first tested. The body weight losses due to the progressive increases in the prerequisite efforts to access food (and exercise) were weak (6-7 %; Fig. 1B), far below our ethics-driven exclusion criteria (- 20%). As opposed to feeding motivation (Fig. 1C), running motivation proved highly sensitive to price/FR schedule increases (Fig. 1D). Females were more motivated for running than males at low prices, a difference however accompanied by a hypersensitivity to price increases (Fig. 1D). Thus, females preferred exercise over food throughout FR1 and FR3 schedules before dropping to an indifference (50%) level (yet reached by males exposed to FR1 schedules; Fig. 1E). Confirming a previous observation (*22*), mice placed under FR1 and FR3 reinforcement schedules without negative impacts on body weights displayed lower daily food intakes (3.91 ± 0.13 g and 3.34 ± 0.40 g respectively in females) than the daily amounts of food (over)eaten by C57Bl/6N females fed *ad libitum* in home cages (4.40 ± 0.11 g, n = 12; p = 0.0076 and p = 0.01 by Mann-Whitney tests following a Kruskal-Wallis analysis of variance). We next verified that our setting did not disrupt the normal daily cycles of feeding and running activities; indeed, maximal levels of these activities, which were sex-dependent for running, were always reached during the dark periods (Fig. 1, F and G). The running duration per (rewarded) sequence, i.e. the time spent running (“consuming” the wheel reward) after paying the price to access the wheel for 1 min, was insensitive to price increases and mouse sex (Fig. 1H). Thus, as opposed to motivation for the wheel, its “consumption” was similar in both sexes. The respective demand elasticity curves for either reward (Fig. 1I) led to higher EVs for food than for exercise and higher exercise EVs in females than in males (Fig. 1J). Analyses of P_max_ values, the maximal prices mice accepted to pay to maintain motivation for each reward (*19*), confirmed EV analyses, except that males did not differ from females (Fig. 1K). On the other hand, Q_0_ values, which are the theoretical drive values for each reward if it was freely accessible (i.e. FR0), displayed a sex difference for exercise, females showing the highest values (Fig. 1L).

**Fig. 1.**
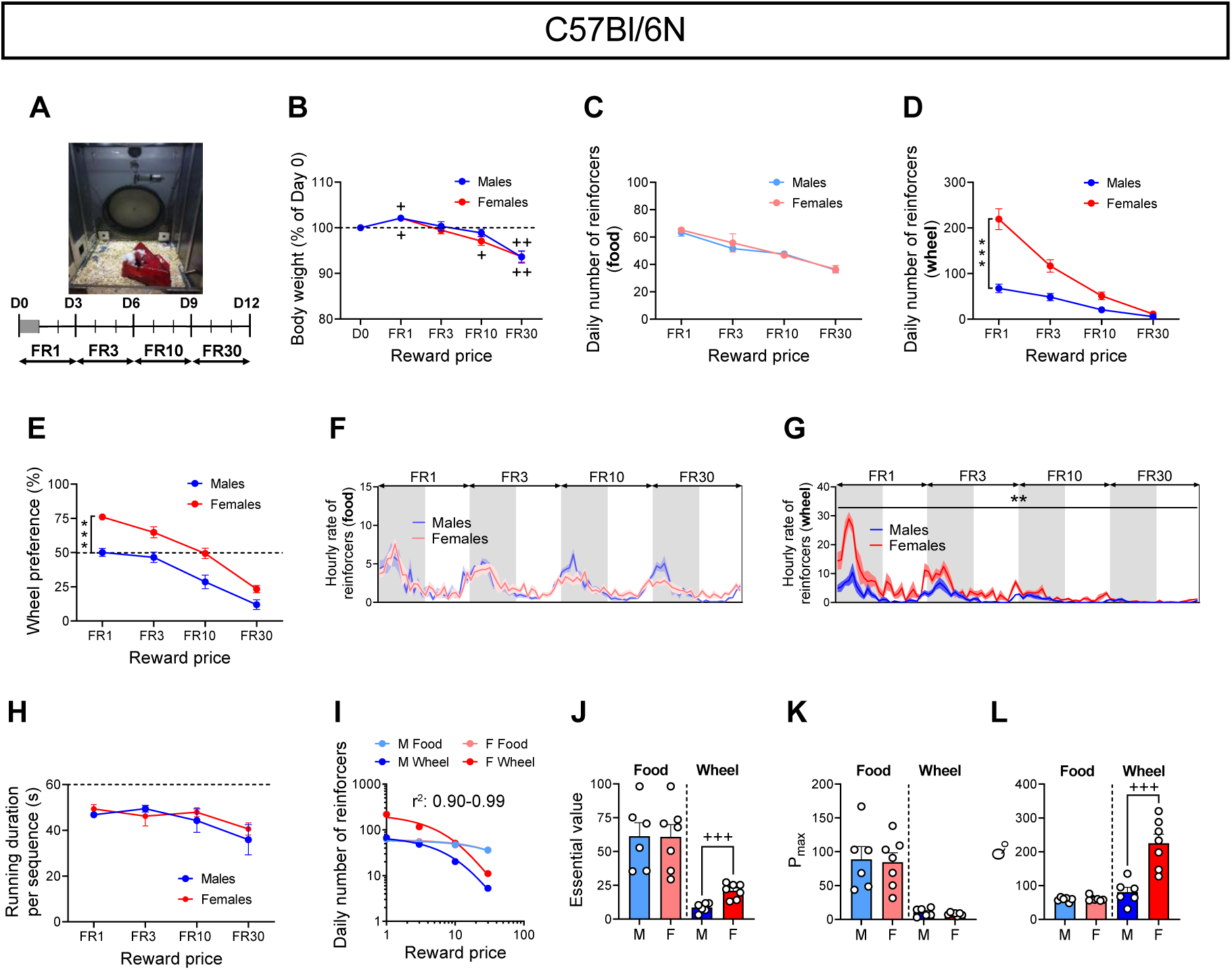
Demand elasticities for food and exercise. (**A**) Operant chamber set-up with active/inactive nose poke ports for food (left) and wheel running (front), unlimited access to a water tip (right) and a shelter filled with nesting material. Each reward was conditioned by fixed ratio (FR) reinforcement schedules of increasing intensities for 12 days. (**B**) Percent body weight changes of male (n = 6) and female (n = 7) C57Bl/6N mice throughout the entire protocol. (**C, D**) Daily numbers of food and exercise reinforcers for each FR schedule. (**E**) Wheel preference over feeding. (**F, G**) Hourly rates of food and exercise reinforcers during the dark/light cycle (dark periods in grey) for each FR schedule. (**H**) Running duration (maximum: 60 sec) for each rewarded sequence of wheel access. (**I**) Demand elasticity curves for food and exercise. (**J**) Food and exercise essential values. (**K, L**) P_max_ and Q_0_ values for food and exercise. All data represent mean ± SEM; **p < 0.01, ***p < 0.001 and ^+^ p < 0.05, ^++^p < 0.01, ^+++^p < 0.001. M and F stand for males and females, respectively. One-tailed Student’s t-test against 100% (B), two-tailed Student’s t-test (J), (K), (L), two-way repeated ANOVA (C), (D), (E), (F), (G), (H).

The 12-day isolation of individual mice in the chambers might be considered stressful, thereby affecting food and exercise drives. To control for this potential confounder, male C57Bl/6N mice were housed either in couples or singly before and during the operant training sessions (fig. S1A). We next compared collectively- and singly-housed mice in the closed economy paradigm under FR1 reinforcement schedules (the price at which food and exercise drives were the highest). The amplitudes of body weight changes (fig. S1B), the respective numbers of reinforcers (fig. S1, C and D), and the wheel preference over food (fig. S1E) did not vary between collectively- and singly-housed mice, suggesting a weak impact of isolation stress.

Having proven the ability of our paradigm to measure the trade-off between feeding and exercise motivations under closed economy conditions, we next aimed at unraveling the mechanisms governing cost (effort)-benefit (value) preference between the two reinforcers.

### Cannabinoid type-1 receptors (CB_1_Rs) exert a positive and selective control on exercise EV through 2-arachidonoylglycerol

The endocannabinoid system (ECS) affects feeding motivation under specific situations (*23–25*). Because it also controls acute exercise motivation during single PR sessions (*12*), we first asked whether this control extends to our closed economy conditions. We thus measured feeding and exercise EVs in male/female CB1R mutants (CB_1_-KO) and their wild-type (CB_1_-WT) littermates (*26*). Except for male WT mice, the closed economy paradigm slightly decreased body weights, females being more sensitive than males (Fig. 2A). In both males (Fig. 2B) and females (Fig. 2C), CB_1_R deletion markedly decreased exercise motivation responses to effort/price demand increases, these changes being associated to a slight sex-dependent decrease in food motivation in male mutants (Fig. 2B). The marked negative impact of CB_1_R deletion on exercise motivation (Fig. 2, B and C) translated into a large drop in the exercise preference ratio over feeding (Fig. 2D). Compared to their WT littermates, male and female mutants ran less during each wheel access consecutive to correct nose poke completion (Fig. 2E). The respective elasticity demand curves (fig. S2A) indicated that feeding EV value did not differ between genotypes (Fig. 2F) whilst running EV, P_max_ and Q_0_ values were decreased in the mutants (Fig. 2, G-I). Exercise EVs decreased by 93 ± 1 % and 90 ± 2 % in male and female CB_1_-KO, compared to their respective WT littermates, indicating that CB_1_Rs play a major role in the chronic control of exercise motivation and demand. Because CB_1_Rs are located at plasma membranes and/or associated to mitochondrial membranes (*27*, *28*), we next investigated the subcellular location of the CB_1_R population controlling exercise motivation. We thus used a knock-in mutant mouse line (called DN22-CB_1_) lacking (DN22-CB_1_-KI) or not (WT) mitochondria-associated CB_1_Rs (*28*). As compared to their WT littermates, male DN22-CB_1_-KI mice displayed no alterations in the reward drives (fig. S3, A and B), in running preference (fig. S3C), in running duration per rewarded sequence (fig. S3D), and in feeding and running EVs (fig. S3, E and F; fig. S2B). These results indicate that the CB_1_R population controlling exercise demand is not located at mitochondrial membranes but likely at plasma membranes.

**Fig. 2.**
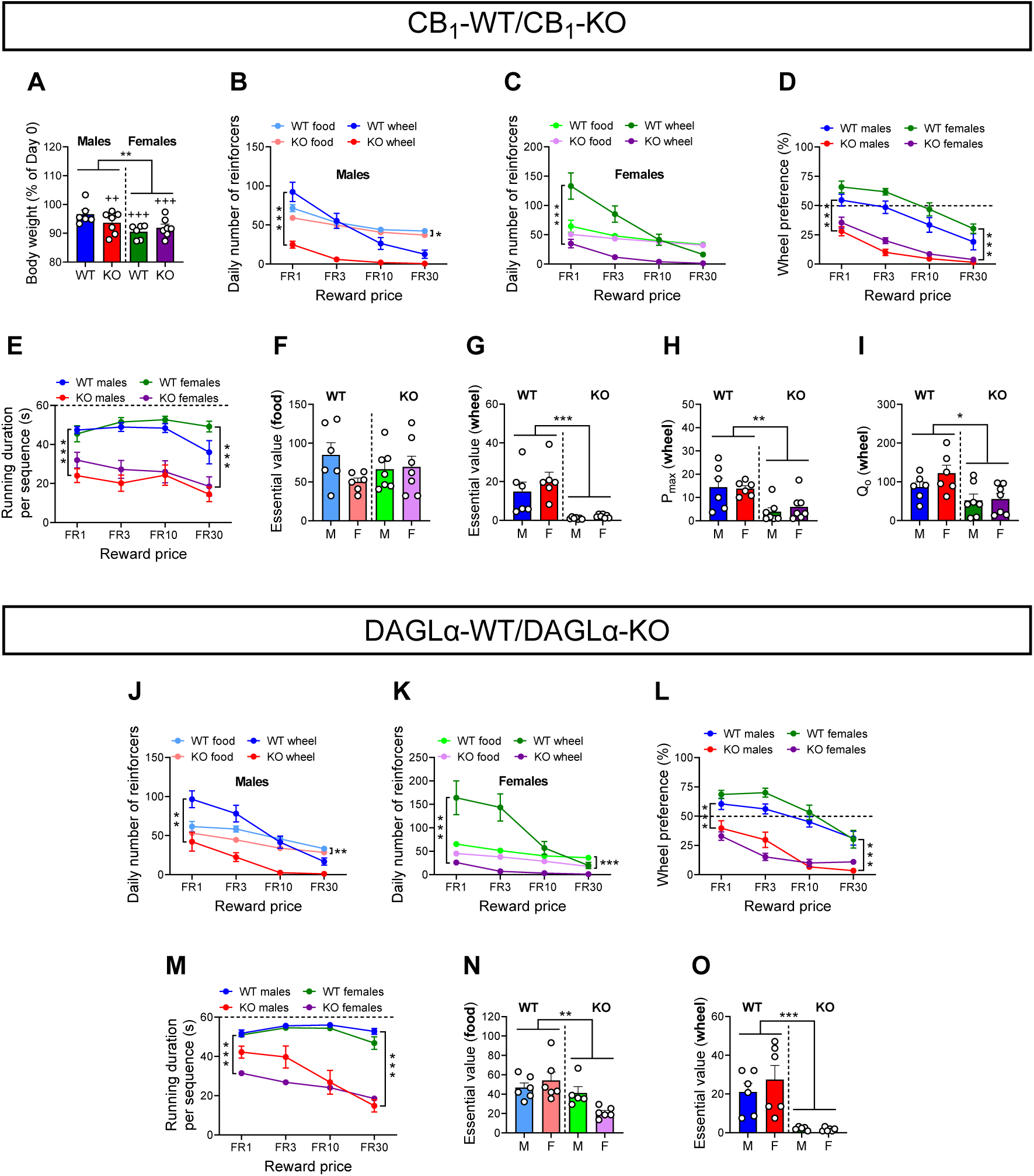
CB_1_ receptors control exercise demand through 2-AG synthesis. (**A**) Percent body weight changes of male (n = 6) and female (n = 6) CB_1_-WT mice and male (n = 7) and female (n = 7) CB_1_-KO mice throughout the entire protocol. (**B**) Daily numbers of food and exercise reinforcers achieved by CB_1_-WT and CB_1_-KO males for each FR schedule. (**C**) Daily numbers of food and exercise reinforcers achieved by CB_1_-WT and CB_1_-KO females for each FR schedule. (**D**) Wheel preference over feeding. (**E**) Running duration for each rewarded sequence of wheel access. (**F**) Food essential values in CB_1_-WT and CB_1_-KO mice. (**G**) Exercise essential values in CB_1_-WT and CB_1_-KO mice. (**H, I**) P_max_ and Q_0_ values for exercise in CB_1_-WT and CB_1_-KO mice. (**J**) Daily numbers of food and exercise reinforcers achieved by n = 6 DAGLα-WT and n = 5 DAGLα-KO males for each FR schedule. (**K**) Daily numbers of food and exercise reinforcers achieved by n = 6 DAGLα-WT and n = 6 DAGLα-KO females for each FR schedule. (**L**) Wheel preference over feeding. (**M**) Running duration for each rewarded sequence of wheel access. (**N, O**) Food and exercise essential values in DAGLα-WT and DAGLα-KO mice. All data represent mean ± SEM; *p < 0.05, **p < 0.01, ***p < 0.001 and ^++^p < 0.01, ^+++^p < 0.001. M and F stand for males and females, respectively. One-tailed Student’s t-test against 100% (A), two-way ANOVA (A), (F), (G), (H), (I), (N), (O), two-way repeated ANOVA (B), (C), (D), (E), (J), (K), (L), (M).

There is evidence for 2-arachidonylglycerol (2-AG) being the preferential endocannabinoid involved in the modulation of reward processes by the ECS (*29*, *30*) (but see (*24*, *31*)). We thus investigated the role of 2-AG in decision-making between food and exercise, doing so through two approaches. The first used acute pharmacology whilst the second one relied on the chronic deletion of the gene encoding for diacylglycerol lipase-alpha (*Dagla*, referred to below as DAGLα), one of the main enzymes involved in 2-AG synthesis (*32*). CB_1_-flox males were first exposed to 4 days of closed economy under a FR1 reinforcement schedule before being injected at the onset of the 4^th^ dark period of the light/dark cycle with a low (30 mg/kg i.p), dose of the 2-AG synthesis inhibitor DO34 (*33*) (fig. S3G). Administration of DO34 almost abolished the drive for exercise (fig. S3I) but it also decreased that for food (fig. S3H). Because feeding motivation was unaltered in CB_1_-KO mice (see above), off-target effects of DO34 (*33*) and/or compensatory mechanisms in CB1-KO animals led us to shift to a selective genetic approach through the use of DAGLα-WT and DAGLα-KO (*34*) male and female mice (fig. S3, J and K). As shown in Fig. 2, DAGLα mutants behaved almost similarly to CB_1_-KO mice in both sexes with regard to exercise. Thus, in males (Fig. 2J) and females (Fig. 2K), the deletion of DAGLα negatively impacted exercise motivation responses to price increases. These impacts of DAGLα deletion extended to exercise preference (Fig. 2L) and exercise duration per rewarded sequence (Fig. 2M). In line with these observations, exercise demand elasticity curves (fig. S2D) led to lower exercise EVs in mutants than in WT mice (Fig. 2O). These genotypic differences were associated to slight decreases in the price-dependent demands for food in DAGLα-KO mice (Fig. 2, J and K; fig. S2D). Thus, DAGLα-KO mice displayed lower food EVs than their WT counterparts (Fig. 2N). However, these EV reductions were markedly less important than those measured for exercise EVs, suggesting that synthesis of 2-AG (and then its release) is a prerequisite for the CB_1_R-mediated control of exercise motivation/demand.

### CB_1_Rs expressed by GABAergic neurons gate exercise demand

As indicated above, the use of Dlx5/6-Cre mice to conditionally delete CB_1_Rs from GABAergic neurons (*35–37*) (GABA-CB_1_-KO mice) has indicated that this CB_1_R subpopulation controls exercise motivation in male mice, as assessed by single PR tests (*12*). Whether this control extends to permanent closed economy conditions and is sex-dependent are issues we assessed using male and female GABA-CB_1_-KO and GABA-CB_1_-WT mice. Exposure to the closed economy protocol promoted a weak body weight loss in GABA-CB_1_-WT mice, but not in the mutants (Fig. 3A). These genotypic differences were not accounted for by differences in feeding motivation (Fig. 3, B and C) but possibly by a reduced motivation for exercise in the mutants (Fig. 3, B and C). As for WT animals, running motivation of male/female mutants was only observed during the dark periods (Fig. 3D), hence excluding mutation consequences on the circadian rhythm. Wheel preference (Fig. 3E) and running performance at each rewarded sequence (Fig. 3F) were both decreased in mutant mice. In contrast to price-dependent demands for food and hence food EV (Fig. 3G; fig. S2E), running EV, P_max_ and Q_0_ values were all diminished in GABA-CB_1_-KO mice, compared to their WT littermates (Fig. 3, H-J; fig. S2E). These data indicate that CB_1_Rs in GABAergic neurons play a significant role in the control of exercise motivation and demand. Moreover, the impacts of CB_1_R deletion on exercise EV in the GABA-CB_1_ line (83-87 % reductions) were close to those measured in CB_1_-KO mice (see above), suggesting that the control of exercise demand by the ECS is mostly mediated by CB_1_Rs located on GABAergic neurons in both males and females.

**Fig. 3.**
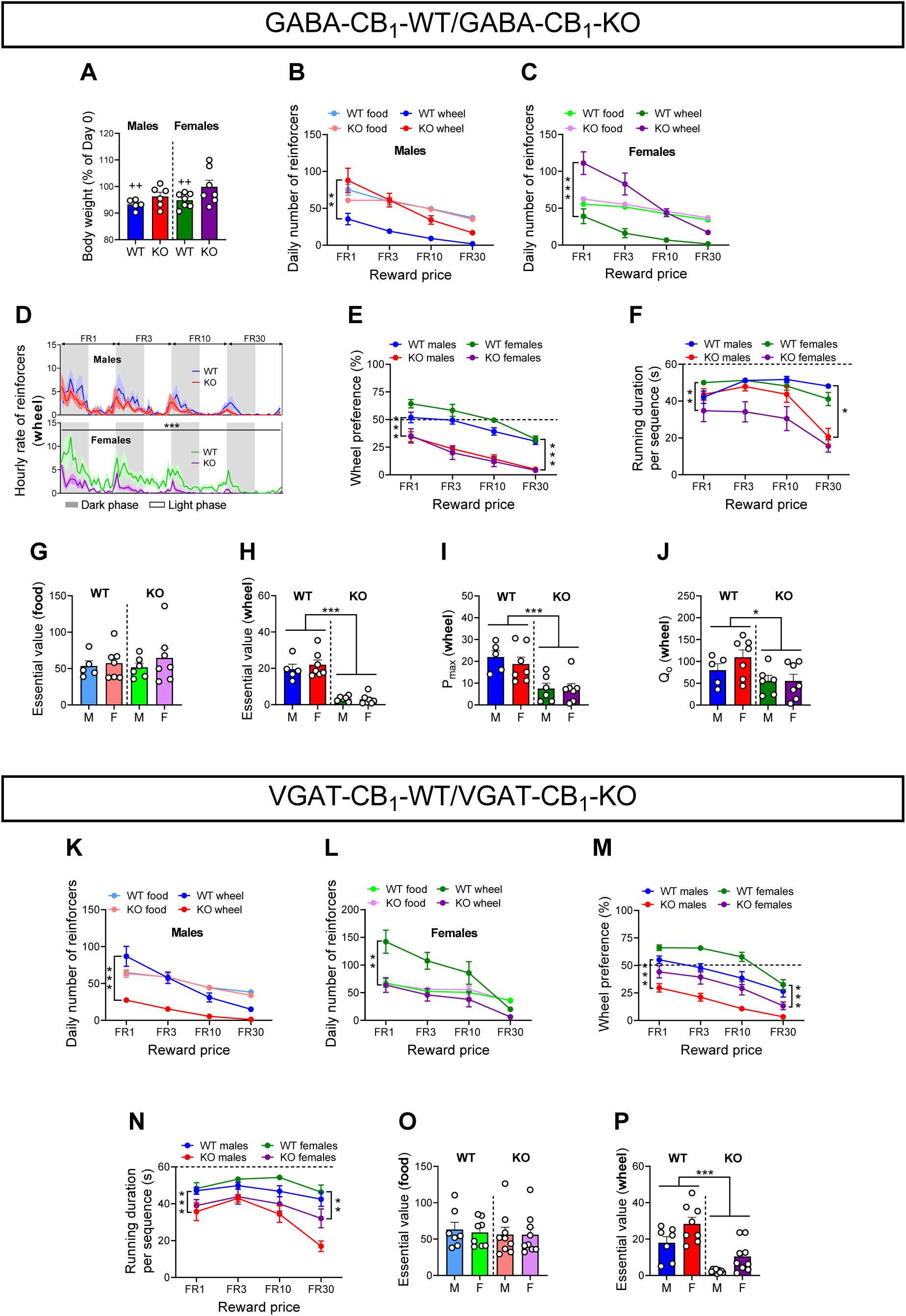
CB_1_ receptors expressed by GABAergic neurons control exercise demand. (**A**) Percent body weight changes of male (n = 5) and female (n = 7) GABA-CB_1_-WT mice and male (n = 6) and female (n = 7) GABA-CB_1_-KO mice throughout the entire protocol. (**B**) Daily numbers of food and exercise reinforcers achieved by male GABA-CB_1_-WT and GABA-CB_1_-KO mice for each FR schedule. (**C**) Daily numbers of food and exercise reinforcers achieved by GABA-CB_1_-WT (n = 7) and GABA-CB_1_-KO (n = 7) females for each FR schedule. (**D**) Hourly rates of exercise reinforcers achieved by male (top) and female (bottom) GABA-CB_1_-WT and GABA-CB_1_-KO mice during the dark/light cycle for each FR schedule. (**E**) Wheel preference over feeding. (**F**) Running duration for each rewarded sequence of wheel access. (**G, H**) Food and exercise essential values in GABA-CB_1_-WT and GABA-CB_1_-KO mice. (**I, J**) P_max_ and Q_0_ values for exercise in GABA-CB_1_-WT and GABA-CB_1_-KO mice. (**K**) Daily numbers of food and exercise reinforcers achieved by VGAT-CB_1_-WT (n = 7) and VGAT-CB_1_-KO (n = 9) males for each FR. (**L**) Daily numbers of food and exercise reinforcers achieved by female VGAT-CB_1_-WT (n = 8) and VGAT-CB_1_-KO (n = 9) mice for each FR schedule. (**M**) Wheel preference over feeding in VGAT-CB_1_-WT and VGAT-CB_1_-KO mice. (**N**) Running duration for each rewarded sequence of wheel access by male and female VGAT-CB_1_-WT and VGAT-CB_1_-KO mice. (**O**) Food essential values in male and female VGAT-CB_1_-WT and VGAT-CB_1_-KO mice. (**P**) Exercise essential values in VGAT-CB_1_-WT and VGAT-CB_1_-KO mice. All data represent mean ± SEM; *p < 0.05, **p < 0.01, ***p < 0.001. M and F stand for males and females, respectively. One-tailed Student’s t-test against 100% (A), two-way ANOVA (A), (G), (H), (I), (J), (O), (P), two-way repeated ANOVA (B), (C), (D), (E), (F), (K), (L), (M), (N).

Dopaminergic neurons express *Dlx5/6* in the arcuate nucleus(*38*), a region involved in the central control of energy balance (*7*). To confirm that the above observations were solely accounted for by CB_1_Rs in GABAergic neurons, we next generated a new double mutant line in which the deletion of the CB_1_-floxed gene is mediated by a Cre recombinase expressed under the control of the promoter of the vesicular GABA transporter (VGAT; (*39*)). After ensuring that the Vgat-Cre recombinase did not bear intrinsic consequences on acute running motivation and performance (fig. S4, A-C), we next crossed CB_1_-floxed mice with Vgat-Cre mice to generate VGAT-CB_1_-WT and VGAT-CB_1_-KO littermate mice. After verifying that CB_1_R were selectively absent from mutant GABAergic cells (e.g. in the striatum and the hippocampus; fig. S4, D and E), we exposed male and female VGAT-CB_1_-WT and VGAT-CB_1_-KO mice to the closed economy paradigm. As for the GABA-CB_1_ line, mutants from the VGAT-CB_1_ line displayed major deficits in exercise, but not feeding, motivation (Figure 3, K and L). These alterations were associated to decreases in wheel preference indices (Fig. 3M), in running duration per sequence (Fig. 3N) and in exercise, but not food (Fig. 3O; fig. S2F), EVs in both sexes (Fig. 3P; fig. S2F).

Mice were only proposed a standard diet in the closed economy setting, raising the possibility that CB_1_Rs in GABAergic neurons exert different controls over exercise and food drives if exposed to a palatable diet (*40*). Indeed, GABA-CB_1_-KO males offered the choice between a sucrose (isocaloric) diet and wheel running fully behaved as the mutants offered a standard diet (fig. S4, F-K), indicating that the control of exercise demand by CB_1_R-expressing GABAergic neurons is food diet-independent.

### Sufficiency of CB_1_Rs expressed by GABAergic neurons for exercise demand

We next tested whether CB_1_Rs on GABAergic neurons were not only necessary but also sufficient for the control of exercise motivation and demand. We thus compared male/female mice bearing a loxP-flanked Stop cassette placed before the open reading frame of the CB_1_ receptor gene (to dampen the expression of CB_1_Rs; STOP-CB_1_ mice) with STOP-CB_1_ mice additionally expressing the Dlx5/6-Cre recombinase so as to re-express CB_1_ receptors selectively in GABAergic neurons (*12*, *41*) (thereafter referred to as GABA-CB_1_-Rescue mice). Whilst these mice did not differ from STOP-CB_1_ mice with regard to feeding motivation and demand (Fig. 4, A, B and F; fig. S2G), exercise-related variables all differed between genotypes. Thus, the weak scores of male and female STOP-CB_1_ mice in running motivation and wheel preference were increased following the selective re-expression of CB_1_Rs in GABAergic neurons (Fig. 4, A-C). Besides, a trend for an increased running duration per rewarded sequence in GABA-CB_1_-Rescue mice could also be observed (Fig. 4D). These increases mostly took place during the dark periods of the light/dark cycles (Fig. 4E). The demand elasticity curves for exercise (fig. S2G) indicated that EV and Q_0_ values, but not that for P_max_, were increased in male and female GABA-CB_1_-Rescue mice, compared to their STOP-CB_1_ littermates (Fig. 4, G-I). These data suggest that CB_1_Rs on GABAergic neurons play a sufficient role on exercise motivation; however, the observation that exercise-related scores of GABA-CB_1_-Rescue mice did not fully reach those measured in GABA-CB_1_-WT mice might question this hypothesis. Although exercise drives in STOP-CB_1_ mice and GABA-CB_1_-KO mice cannot be compared, it is noteworthy that male and female STOP-CB_1_ mice displayed 74 % and 78% decreases in exercise EVs, compared to their respective GABA-CB_1_-Rescue littermates (Fig. 4G). Actually, these percent changes were similar to those measured in male and female GABA-CB_1_-KO mice (see above).

**Fig. 4.**
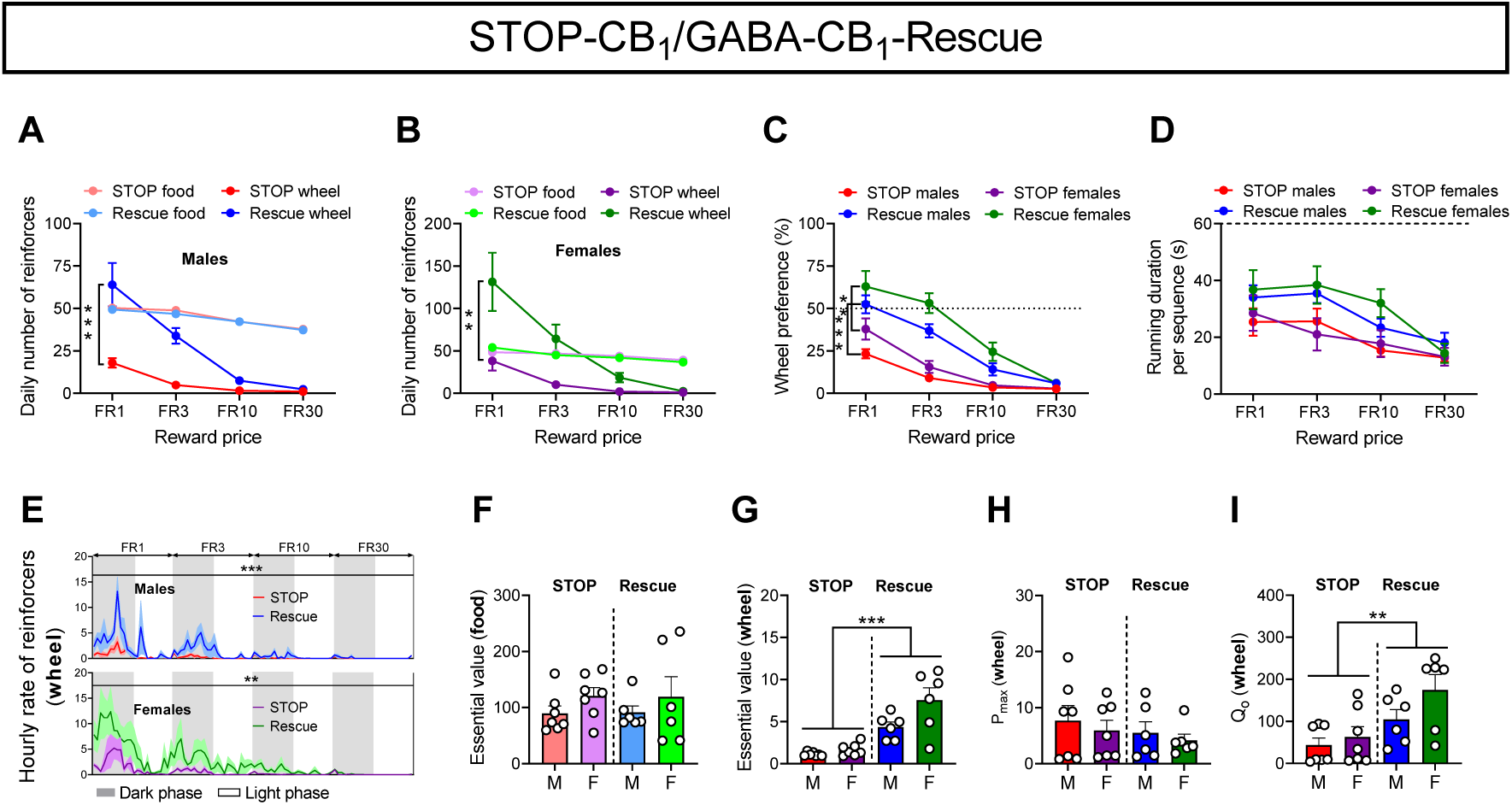
CB_1_ receptors on GABAergic neurons are necessary for the control of exercise demand. (**A**) Daily numbers of food and exercise reinforcers achieved by male STOP-CB_1_ (n = 7) and GABA-CB_1_-Rescue (n = 6) mice. (**B**) Daily numbers of food and exercise reinforcers achieved by female STOP-CB_1_ (n = 7) and GABA-CB_1_-Rescue (n = 6) mice. (**C**) Wheel preference over feeding. (**D**) Running duration for each rewarded sequence of wheel access. (**E**) Hourly rates of exercise reinforcers achieved by male (top) and female (bottom) STOP-CB_1_ and GABA-CB_1_-Rescue mice during the dark/light cycle for each FR schedule. (**F**) Food essential values in male and female STOP-CB_1_ and GABA-CB_1_-Rescue mice. (**G-I**) Exercise essential values, and P_max_ and Q_0_ values in STOP-CB_1_ and GABA-CB_1_-Rescue mice. All data represent mean ± SEM; **p < 0.01, ***p < 0.001. M and F stand for males and females, respectively. Two-way ANOVA (F), (G), (H), (I), two-way repeated ANOVA (A), (B), (C), (D), (E).

### Roles of astrocytic and glutamatergic CB_1_R subpopulations on the respective demands for food and exercise

Although the above data assign to CB_1_Rs on GABAergic neurons a major role in the control of exercise motivation, other CB_1_R-expressing cells, placed upstream of these GABAergic neurons, might also participate in this control. Indeed, CB_1_R-expressing astrocytes and glutamatergic neurons have major influences on reward processes, especially in the ventral striatum (*42–44*). We thus addressed this issue using appropriate conditional CB_1_R mutants. Male mice lacking CB_1_Rs in astrocytes (GFAP-CB_1_-KO mice; (*45*, *46*)) did not behave differently from their WT (GFAP-CB_1_-WT) littermates with respect to feeding- and exercise-related motivation, demand and performance variables (fig. S5, A-D; fig. S2H). On the other hand, the deletion of CB_1_Rs in glutamatergic neurons in male mice (termed thereafter Glu-CB_1_-KO; (*12*, *35*, *37*)) reduced running motivation and the wheel preference ratio (fig. S5, E-G). Furthermore, demand curves (fig. S2I) revealed that besides a 63% reduction in the exercise EV, mutants also displayed a significant decrease in feeding EV (fig. S5H). We next wondered whether selectively re-expressing CB_1_Rs in glutamatergic neurons (Glu-CB_1_-Rescue; (*47*)) of mice lacking CB_1_R expression (i.e. STOP-CB_1_ mice; see above) would rescue running and food demands. In sharp contrast with the rescue of CB_1_Rs in GABAergic neurons, re-expression of CB_1_Rs in glutamatergic neurons of male STOP-CB_1_ mice proved inefficient in rescuing exercise- (and feeding-) dependent variables (fig. S5, I-L; fig. S2J).

Taken together, these results indicate that CB_1_Rs on glutamatergic neurons are involved in the respective controls of food and exercise demands. However, the control exerted by this population on exercise variables is (partly) necessary but not sufficient. We thus prioritized the quest for the location of the CB_1_R-expressing GABAergic neuron sub-population bearing both necessary and sufficient roles on exercise motivation and demand.

### Ventromedial striatal GABAergic neurons selectively gate exercise demand

The observation that an adeno-associated virus (AAV)-mediated deletion of CB_1_Rs from the VTA of CB_1_-floxed males did not affect food and exercise seekings (data not shown) led us to shift to another major hub in reward motivation, i.e. the nucleus accumbens (NAc) (*48*). We first focused on the medial part of the ventral striatum (ventromedial striatum; VMS) owing to its regulation of reward-guided behaviors (*49*, *50*). Thus, we bilaterally injected an AAV expressing a CAG-Cre-GFP construct (or a control AAV expressing a CAG-hrGFP construct) into the VMS of male and female CB_1_-floxed mice (Fig. 5, A and B). The conditional deletion of VMS CB_1_R in males and females altered neither feeding motivation (Fig. 5, C and E) nor food EV (Fig. 5K; fig. S2L). On the other hand, exercise motivation, whether examined on daily (Fig. 5D) or hourly (Fig. 5G) bases, was decreased in males but not in females (Fig. 5, F and H). This sex-dependent response included the exercise preference over feeding (Fig. 5I) and the time spent running per rewarded sequence (Fig. 5J). Analyses of exercise demand elasticities confirmed these latter results (Fig. 5L; fig. S2L). For the sake of specificity, we next injected the same viruses into another reward-associated ventrostriatal region, namely the ventrolateral striatum (*49*, *50*) (VLS; fig. S6A). Indeed, CB_1_R deletion in the VLS of CB_1_R-floxed male mice did not affect exercise-related variables or food motivation and demand (fig. S6, B-E; fig. S2K).

**Fig. 5.**
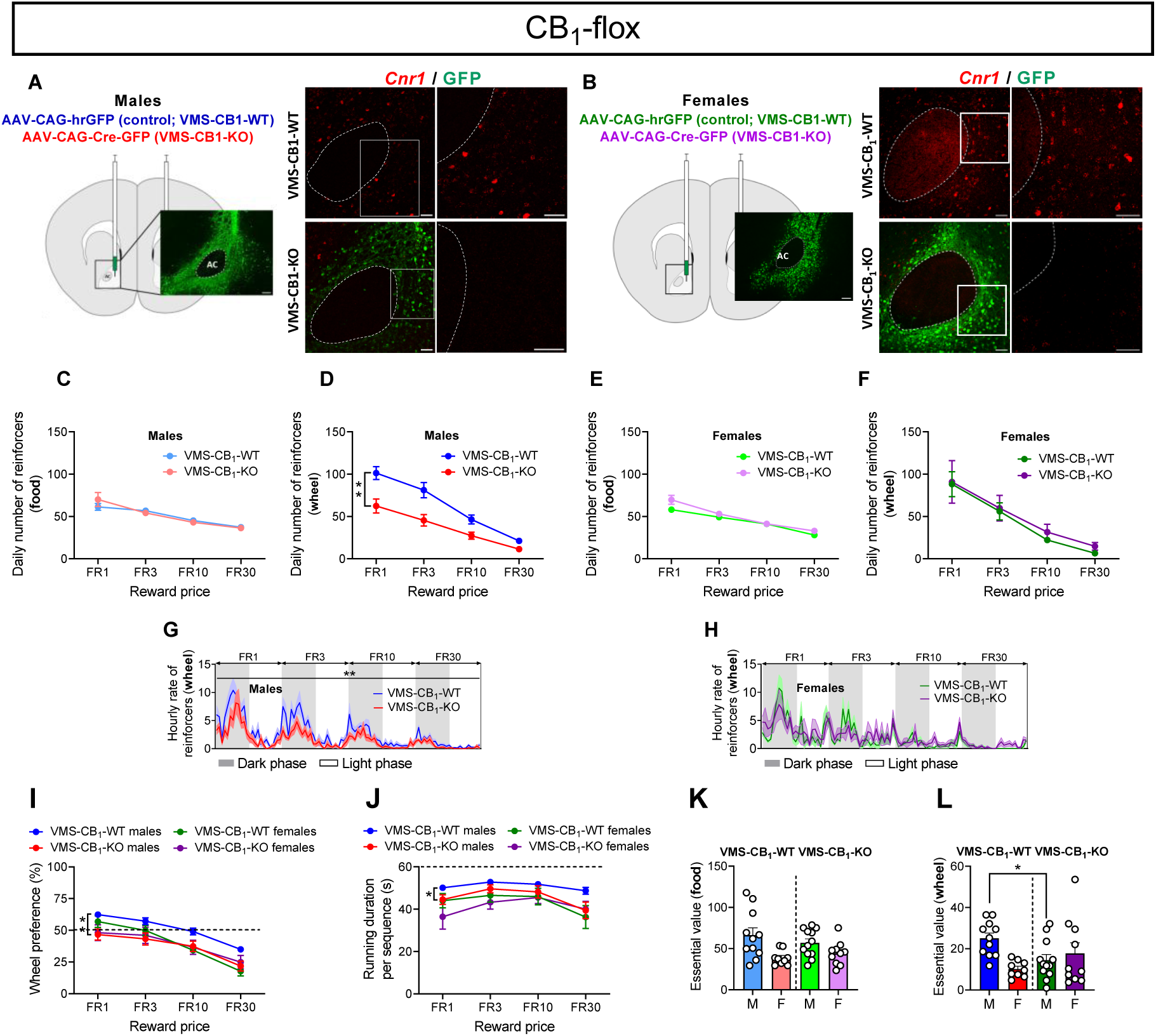
Ventromedial striatal CB_1_ receptor-expressing GABAergic neurons gate exercise demand in males, but not in females. (**A, B**) Viral-based strategies to delete CB_1_ receptors from the ventromedial striatum of male and female CB_1_-floxed mice with representative images of GFP immunostaining (left; scale bar = 100 µm) and detailed fluorescence *in situ* hybridization for *Cnr1* transcripts and GFP immunostainings (right; scale bars = 50 µm). (**C, D**) Daily numbers of food and exercise reinforcers achieved by male mice injected with the inactive virus (VMS-CB_1_-WT, n = 11) and the active virus (VMS-CB_1_-KO, n = 12) for each FR schedule. (**E, F**) Daily numbers of food and exercise reinforcers achieved by female VMS-CB_1_-WT (n = 9) and VMS-CB_1_-KO (n = 10) mice for each FR schedule. (**G, H**) Hourly rates of exercise reinforcers achieved by male and female VMS-CB_1_-WT and VMS-CB_1_-KO mice during the dark/light cycle for each FR schedule. (**I**) Wheel preference over feeding in VMS-CB_1_-WT and VMS-CB_1_-KO mice. (**J**) Running duration for each rewarded sequence of wheel access by VMS-CB_1_-WT and VMS-CB_1_-KO mice. (**K, L**) Food and exercise essential values in VMS-CB_1_-WT and VMS-CB_1_-KO mice. Two-way ANOVA (K), two-way repeated ANOVA (B), (C), (D), (E), (F), (G), (H), (I), (J), Kruskal-Wallis ANOVA (L).

As VMS CB_1_Rs are necessary for exercise motivation and demand in male mice, we next investigated whether this receptor subpopulation is expressed by GABAergic neurons. We injected an AAV expressing a CAG-flex-CB_1_-GFP construct (or a CAG-flex control virus) in the VMS of GABA-CB_1_-KO male mice (Fig. 6A). This manipulation, which led to the selective re-expression (rescue) of CB_1_Rs in VMS GABAergic neurons (Fig. 6A), did not affect food motivation and demand elasticity (Fig. 6B, F; fig. S2M) but selectively increased exercise motivation (Fig. 6, B and C), wheel preference (Fig. 6D), running duration per rewarded sequence (especially at the highest price; Fig. 6E), and running EV (Fig. 6G; fig. S2M). These results indicated that CB_1_Rs located on VMS GABAergic neurons play necessary and sufficient roles on value-based decision making for exercise in male mice. Taken together, these observations thus mirrored the data obtained respectively in male GABA-CB_1_-KO mice and in GABA-CB_1_-Rescue mice, compared to their respective littermate controls (see above). However, the observation that VMS GABAergic neurons are not necessary for exercise motivation in females did not exclude the possibility that these neurons play a sufficient role. This hypothesis was tested in GABA-CB_1_-KO females using the same viral approaches as those used in males (fig. S7A). The results indicate that CB_1_Rs in VMS GABAergic neurons are neither necessary (see above; Fig. 5) nor sufficient for exercise (and food) motivation and demand (fig. S7, B-E; fig. S2M). Thus, although CB_1_R-expressing GABAergic neurons gate in a sex-independent manner the drive for exercise under increasing effort requirements, their brain location is sex-specific.

**Fig. 6.**
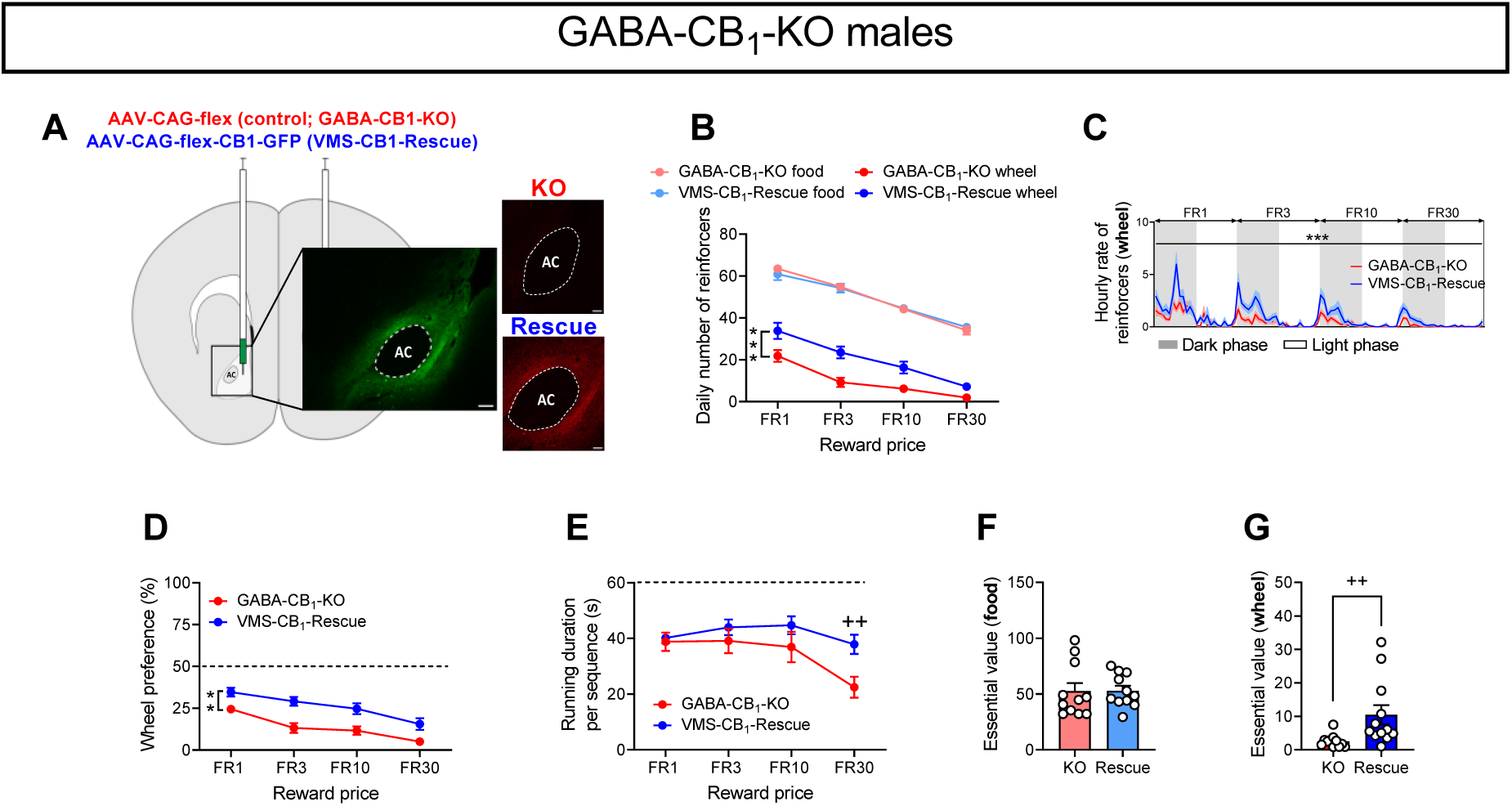
Ventromedial striatal re-expression of CB_1_ receptors in GABAergic neurons of GABA-CB1-KO males increases exercise demand in males. (**A**) Viral-based strategies to re-express CB_1_ receptors in the ventromedial striatum of GABA-CB_1_-KO male mice with representative images of GFP immunostainings (scale bar = 100 µm) and CB_1_ receptor immunostainings (scale bars = 50 µm). (**B**) Daily numbers of food and exercise reinforcers achieved by male mice injected with the inactive virus (GABA-CB_1_-KO, n = 11) and the active virus (VMS-CB_1_-Rescue, n = 12) for each FR schedule. (**C**) Hourly rates of exercise reinforcers achieved by male GABA-CB_1_-KO and VMS-CB_1_-Rescue mice during the dark/light cycle for each FR schedule. (**D**) Wheel preference over feeding in male GABA-CB_1_-KO and VMS-CB_1_-Rescue mice. (**E**) Running duration for each rewarded sequence of wheel access by male GABA-CB_1_-KO and VMS-CB_1_-Rescue mice. (**F, G**) Food and exercise essential values in male GABA-CB_1_-KO and VMS-CB_1_-Rescue mice. All data represent mean ± SEM; *p < 0.05, **p < 0.01, ***p < 0.001 and ^++^p < 0.01. M and F stand for males and females, respectively. Mann-Whitney test (F), (G), two-way repeated ANOVA (B), (C), (D), (E).

### Exercise demand is controlled by ventrostriatal CB_1_R-expressing GABAergic interneurons

Ventrostriatal medium spiny neurons (MSNs) are thought not to express CB_1_Rs (*51*, *52*) but a recent study suggested their presence, albeit at extremely low levels, in dopamine (DA) receptor 1 (D1R)-expressing MSNs (*53*). In our hands, CB_1_R mRNA was not detectable in ventrostriatal D1R- (fig. S7A) and DA receptor 2 (D2R)-expressing MSNs (fig. S7H). Because co-expression of D1Rs (and/or D2Rs) and CB_1_Rs might occur below our detection limits, we next used two mouse lines, hereafter called D1-CB_1_ (initially called DrD1-CB_1;_ (*54*)) and A2A-CB_1_, wherein mutants lack CB_1_Rs in striatal D1- and D2-expressing neurons respectively (*54*). The comparison between these mutants and their respective WT littermates revealed no difference in any exercise- or food-related variable under investigation (fig. S8, B-N; fig. S2, N and O). To ensure that the involvement of CB_1_R-expressing MSNs, if any, could be excluded, we then turned to a viral approach allowing the removal of CB1Rs from all VMS principal neurons. Thus, we bilaterally injected an AAV bearing the expression of Cre recombinase under the control of the Ca^2+^/calmodulin-dependent kinase IIα (AAV-CaMK-Cre) promoter in the VMS of male CB_1_-floxed mice (fig. S8O). The results confirmed the data gathered in CB_1_R-expressing MSN mutants (fig. S8, P-U; fig. S2P), suggesting that VMS CB_1_R-expressing GABAergic interneurons, but not MSNs, control exercise motivation.

Evidence for the absence of CB_1_Rs in ventrostriatal cholinergic interneurons (*42*) suggests that CB_1_R-expressing inhibitory neurons are solely accounted for by two populations of fast-spiking interneurons (FSIs) in the ventral striatum (*52*, *55*). Because the first one co-expresses parvalbumin (PV) but not the second one (*52*), we next tested whether the selective deletion of CB_1_Rs from PV-expressing neurons had any consequence on exercise motivation/demand. As a prerequisite, we first examined acute/chronic exercise motivation in male PV-Cre^+^ (PV-Cre) and PV-Cre^-^ (PV-WT) littermates (from male PV-Cre and female C57Bl/6N mouse crossings) to ensure that the recombinase lacked intrinsic impact on exercise motivation. After checking that the number of PV-expressing cells was not altered in PV-Cre mice (e.g. in dorsal striatum: fig. S9A), we next tested these mice under FR1, FR3 and PR reinforcement schedules (fig. S9B). Indeed, no phenotypic difference could be observed (fig. S9, C and D). When PV-Cre mice and PV-WT mice were tested for 3 consecutive days under continuous FR1 reinforcement schedules in the closed economy setting (fig. S9E), neither the numbers of reinforcers (fig. S9, F and G) nor the preference for wheel running and exercise performance (fig. S9, H and I) differed between genotypes. After generating PV-CB_1_-KO and PV-CB_1_-WT littermates - by crossing PV-Cre mice with CB_1_-floxed mice - we observed that the expression of CB_1_R transcripts, which were present in 92.8 % of dorsostriatal PV cells in PV-CB_1_-WT mice was reduced by 50 % in PV-CB_1_-KO mice (fig. S9J). Whether this ratio applies to the NAc is unknown as similar assays therein proved unsuccessful and/or unreliable. Under closed economy settings, the price-elicited decrease in wheel running motivation (fig. S9L) and its corresponding elasticity (fig. S2Q) did not differ between WT and mutant males. These findings extended to other exercise-related variables as well as to feeding motivation (fig. S9, K and M-P), suggesting that VMS CB_1_R-expressing GABAergic interneurons controlling exercise motivation belong to the non-PV FSI subtype.

## DISCUSSION

A recent review on the weak predictive and therapeutic values of most commonly used animal models of psychopathology recommended new paradigms, with longer lasting durations and environmental settings enriched and more naturalistic than the present ones (*16*). The goal of our closed economy model, based on permanent cost-benefit decisions, aimed at approaching these criteria whilst specifically addressing one complex dimension of reward processes, namely motivation (*2*, *56*). As opposed to the neurobiology of feeding motivation which is investigated through operant conditioning paradigms, including in home cages (*22*, *57*), only few studies have focused on the bases of exercise motivation. Indeed, most studies use free running wheels (*15*) although free wheel running performance is not a reliable predictor of running motivation (*12*, *13*, *58*). In addition to specifically addressing exercise motivation, our paradigm allows an objective comparison between feeding and exercise incentive properties which are most often individually addressed (but see: (*12*, *59*)). Being permanent and providing a reward choice, our closed economy protocol thus differs from the preceding ones, which are of short duration and/or address the demand elasticities of a single reward. In the present study, food and exercise demands followed innate light/dark cycles and showed different amplitudes, mice being less sensitive to price increases to get food than exercise owing to the vital need for the former reward. In line with prior evidence that exercise is a strong reward in rodents (*9–12*), mice preferred to exercise than to feed at low prices. Thus, these economic conditions allowed mice to perform extra efforts to exercise in addition to those needed for vital feeding.

Whilst the neurobiological bases of exercise performance (*15*, *36*, *60*) and exercise preference (*10*, *61*), including over another reward (*62*, *63*), are somewhat documented, how (and where) exercise motivation takes place is only partly uncovered. We have proposed that VTA dopaminergic systems are involved (*12*, *13*), a hypothesis recently confirmed (*64*). Within the VTA, there is evidence that CB_1_Rs gate exercise motivation (*12*), likely by modulating GABAergic afferences to dopaminergic neurons. Whether this CB_1_R-mediated control extends to permanent cost-benefit situations is an issue we next addressed. Indeed, the deletion of CB_1_Rs triggered a profound decrease in exercise motivation and demand, as assessed by extremely low exercise EVs. Conversely, feeding demand and EV remained unaltered in CB_1_R mutants, which translated in a decreased wheel preference over food in these animals. The observation that feeding demand was insensitive to CB_1_R deletion is in line with the exostatic role of CB1Rs with respect to feeding (*40*). Indeed, CB_1_Rs gate the need for palatable food (*40*), but not that for a standard diet, unless restricted feeding (*25*) or fasting (*37*) precedes refeeding. Of particular note was the additional finding that CB_1_R deletion decreased the running duration per rewarded sequence. This observation contrasts with our previous report that CB_1_-KO and CB_1_-WT mice did not differ with respect to that variable when daily tested for 1h-sessions (*12*), hence suggesting that under chronic effort-based demands, motivation and consumption cannot be disentangled.

The DAGL inhibitor DO34 decreased the efforts made to access exercise, suggesting that the CB_1_R control of exercise motivation is mediated by 2-AG. In contrast with the behaviors of CB_1_-KO mice, DO34 also lowered food seeking, suggesting that DO34 acted through CB_1_R-independent mechanisms. Indeed, DO34 might target other enzymes than DAGL (*33*), including the postsynaptic 2-AG degradation enzyme α/β-hydrolase domain 6 (ABHD6). Remarkably, ABHD6 deletion in the NAc decreases motivation for palatable food (*24*). Using DAGLα-WT and DAGLα-KO mice, we observed that the respective inhibitory effects of DAGLα deletion on exercise motivation/demand equated those measured in male and female CB_1_-KO mice. These results indicate that 2-AG signaling is crucially involved in the ECS-dependent control of exercise motivation and demand whilst the additional impact of DAGLα deletion on food motivation might be mediated by non-CB_1_Rs (*32*).

Male and female GABA-CB_1_-KO mice behaved as constitutive CB_1_-KO mice with regard to either reward, suggesting that most, if not all, CB_1_Rs controlling exercise EV are expressed by GABAergic neurons. The data gathered in GABA-CB_1_-KO male and female mice were replicated in VGAT-CB_1_-KO mice, thereby confirming that CB_1_R-expressing GABAergic neurons gate exercise motivation/demand. This control seems food diet-independent as similar results were observed in mice offered a sucrose diet. In addition, selectively re-expressing CB_1_R in GABAergic neurons of CB_1_R-silenced male/female mice increased exercise demand and preference over food. Although the latter results indicate that CB_1_R-expressing GABAergic neurons play both necessary and sufficient roles on exercise motivation/demand, we could not exclude the downstream/upstream involvement of other CB_1_R-expressing cell types. Because CB_1_R-expressing glutamatergic neurons and astrocytes play pivotal roles in incentive processes (*30*, *55*), we next focused on these cell types. Whereas deleting CB_1_Rs from astrocytes did not impact mouse behaviors, CB_1_Rs from glutamatergic neurons proved necessary, but not sufficient, for the control of food demand. Although the former finding is at first glance in line with the role of CB_1_R-expressing glutamatergic on food intake, it is noteworthy that such a role was established using either unconditioned fasting/refeeding experiments (*37*) or conditioned feeding responses to a palatable diet (*23*) (but see: (*12*)). Because the decreased motivation for food in Glu-CB_1_-KO mice was not observed in full CB_1_-KO mice, it suggests different compensatory mechanisms in the mutants and/or the presence of a CB_1_R subpopulation playing an opposite role on food demand. These results open future routes of investigation aimed at identifying which NAc-projecting CB_1_R-expressing glutamatergic neurons (*42*, *55*, *65*) are involved in food demand. In addition, CB_1_Rs expressed by glutamatergic neurons were also involved in exercise EV. Of note was the observation that these neurons control exercise EV, hence in contrast with their dispensability in the control of acute exercise motivation (*12*). As opposed to CB_1_R-expressing GABAergic neurons, CB_1_R-expressing glutamatergic neurons proved only necessary for exercise demand, suggesting that the latter population is located upstream of the former one (see below for further discussion).

The key role of NAc DA in effort-based decision-making (*2*, *56*) and its tight direct/indirect regulation by the ECS (*55*) prompted us to focus on that brain region. Moreover, recent works revealed that NAc D1-expressing and D2-expressing MSNs control in a MSN subtype-specific manner acute running motivation and performance (*64*, *66*). Herein, we show that VMS, but not VLS, CB_1_Rs sex-specifically gate exercise motivation/demand. Because ventrostriatal cholinergic interneurons lack CB_1_Rs (*42*), this suggested that MSNs and/or GABAergic interneurons were the exclusive targets of the AAVs used to delete CB_1_Rs. Consistently, the selective re-expression of CB_1_Rs in the VMS of male GABA-CB_1_-KO mice increased to a major extent the drive for exercise. Whether VMS CB_1_R-expressing GABAergic cells are fully responsible for the decreased exercise motivation of GABA-CB_1_-KO males is unknown. Thus, the comparison between the drops in exercise EVs between VMS-CB_1_-KO and GABA-CB_1_-KO males is confounded by the fact that CB_1_Rs are deleted at early developmental stages in GABA-CB_1_-KO mice whilst the AAV-mediated deletion of these receptors occurs at adulthood. Considering the observed sex differences in the involvement of VMS CB_1_Rs in exercise demand, an important issue is the location of the CB_1_R subpopulation controlling exercise EV in females. Sex differences in the mechanisms governing reward processes are documented (*67*), but determining their exact anatomical substrates will prove a long experimental process.

The belief that ventrostriatal MSNs do not express CB_1_Rs (*52*) has been recently challenged by the detection of low levels of *cnr1* transcripts in NAc D1-expressing MSNs (*53*) and functional evidence for CB_1_Rs at NAc D1-expressing MSN terminals in the lateral hypothalamus (*68*). Although we failed to detect CB_1_R expression in NAc MSNs, we could not exclude that CB_1_Rs are expressed at undetectable levels in these cells. However, neither the deletion of CB_1_Rs from D1-expressing or D2-expressing MSNs (*54*) nor injection of a CaMKII-promoter associated AAV to conditionally remove CB_1_Rs from male VMS MSNs altered exercise motivation and demand. These results provided definitive evidence for an unforeseen role of VMS GABAergic interneurons in the control exercise motivation/demand.

One important limitation of our study lies in our inability to further define which GABAergic interneurons are precisely involved in chronic exercise motivation. This limitation is mostly accounted for by technical limits owing to their sparsity in the NAc and hence the limited knowledge on their molecular properties. As mentioned above, NAc CB_1_Rs are expressed by two subpopulations of FSIs which differ by the presence/absence of PV co-expression (*52*). Thus, 65% of PV-expressing cells also co-express CB_1_Rs whilst 50 % of CB_1_R-expressing cells are labelled with PV. Owing to FSI feed-forward properties in the NAc, these cells, which include CB_1_R-expressing cells (*52*, *69*), mediate the motivation and/or the preference for natural rewards and drugs (*70*). Moreover, FSIs might be targeted by 2-AG released from postsynaptic MSNs (*71*, *72*), thus reinforcing the hypothesis that FSIs might control exercise motivation. However, the deletion of CB_1_Rs from PV-expressing cells did not alter exercise motivation and demand in male mice. At first glance, this observation might be accounted for by the incomplete reduction of CB_1_ transcripts in PV-CB_1_-KO mice (at least in the dorsal striatum). This last explanation is however too simplistic considering that the constitutive deletion of CB_1_Rs promotes a developmental loss of PV cells (*73*), and that only half of NAc PV-expressing cells co-express CB_1_Rs. Another possibility is that the CB_1_R-expressing interneurons controlling exercise motivation/demand actually belong to the PV-negative FSIs mentioned above. Because this last cell type has not been fully characterized, tools to confirm this hypothesis are however presently lacking. Despite this uncertainty, a hypothetical circuitry can still be proposed wherein (i) 2-AG released from VMS MSNs would (ii) stimulate CB_1_Rs located on this FSI population, (iii) thereby disinhibiting MSN activity and hence (iv) ultimately activating VTA dopaminergic neurons (Fig. 7). This would also explain why CB_1_R-expressing glutamatergic neurons are necessary, but not sufficient, for an appropriate drive for exercise. Thus, if these neurons belong to the population of CB_1_R-expressing corticolimbic glutamatergic neurons which innervate VMS FSIs (*74*), it is expected that following their stimulation, the increased excitatory tone upon FSIs (resulting from the absence of CB_1_Rs in these excitatory neurons) amplifies FSI feed-forward inhibitory potencies and hence reduce exercise motivation/demand.

**Fig. 7.**
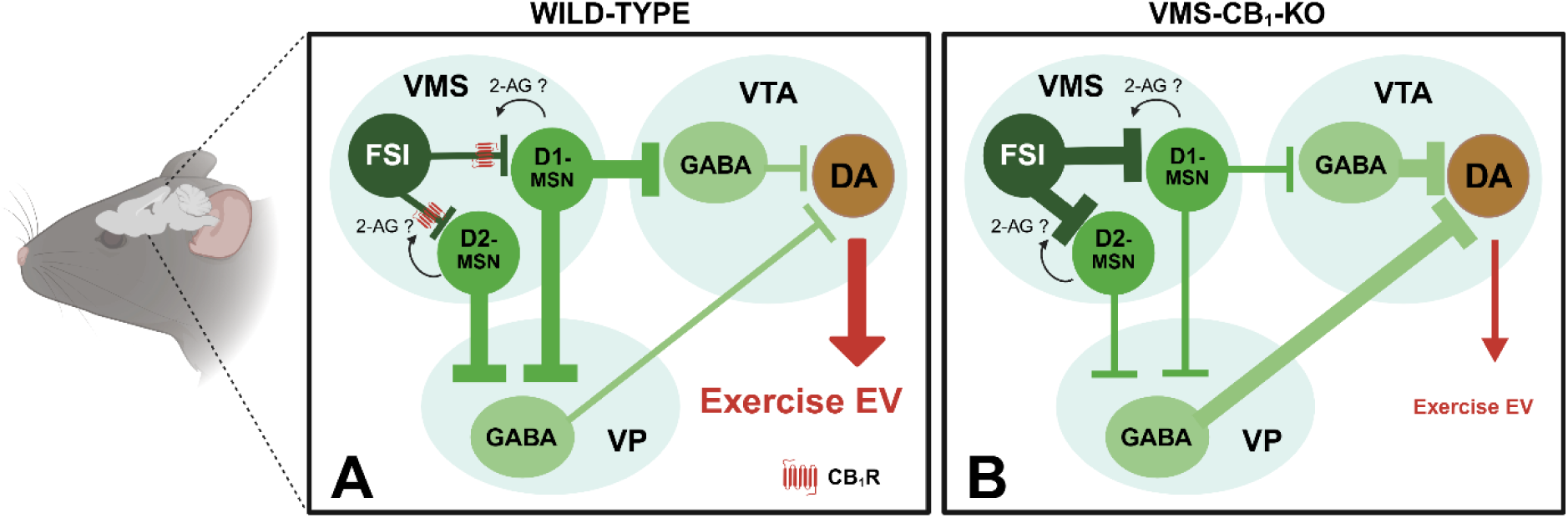
Schematic illustration of how the deletion of CB_1_Rs from ventromedial striatal (VMS) GABAergic interneurons might impair exercise demand (EV) in male mice. (**A**) In control (WT) animals, the release of endocannabinoids (possibly 2-arachidonylglycerol; 2-AG) from D1- and/or D2-expressing medium spiny neurons (MSNs) stimulates CB_1_Rs at presynaptic fast spiking interneurons (FSIs), thereby decreasing their inhibitory input on MSNs. The increased tone of MSNs then reduces ventral pallidum (VP) and/or ventral tegmental area (VTA) GABAergic brakes on VTA dopaminergic neurons, this disinhibition allowing exercise demand to be effectively fulfilled. (**B**) The deletion of CB_1_Rs in the VMS does not permit anymore a FSI-mediated disinhibition of MSNs, hence leaving intact VP and/or VTA GABAergic brakes on VTA dopaminergic neurons. As a consequence, exercise demand is weak and vanishes when strong effort prerequisites are applied. Note that this scheme is not valid for female animals (see Discussion). Created in BioRender. Hurel, I. (2026)

In conclusion, we provide a unique model wherein feeding and exercise drives, and their elasticity responses to increasing efforts, can be permanently assessed. Thereby, we reveal the sex-dependent necessary and sufficient role of VMS CB_1_R-expressing GABAergic interneurons in the control of exercise motivation. Beyond the present study, we believe that the closed economy developed herein holds promising future interests in the quest for the CNS bases of pathological imbalances in food/exercise choices, such as obesity and restricted anorexia nervosa.

## Materials and methods

### Animals

The experiments described above obeyed the French (Décret 2013-118) and European (2010/63/EU) 3R rules on animal experimentation and were approved by the French Ministry of Research (APAFIS # 22435, # 32662, and #47975 to F.C).

This study involved 6-12 week-old male/female C57Bl/6N mice (Elevage Janvier, France) and 8-16 week-old (at the onset of the experiments) male/female constitutive and conditional CB_1_R mutant (KO) and wild-type (WT) animals from our breeding facilities. These animals included (i) floxed-CB_1_R (CB_1_-flox) mice, (ii) CB_1_-KO mice and their CB_1_-WT littermates (*12*, *26*, *35*), (iii) mice lacking (DAGLα-KO) or not (DAGLα-WT) DAGLα (*33*), the synthesis enzyme for 2-AG, (iv) mice lacking (or not) CB_1_Rs from GABAergic neurons by means of either the Dlx5/6-Cre recombinase (*12*, *35*, *37*) (GABA-CB_1_-KO/GABA-CB_1_-WT) or a Cre recombinase driven by the promoter of the vesicular GABA transporter protein (*Vgat* or *Slc32a1*(*39*); VGAT-CB_1_-KO/VGAT-CB_1_-WT), (v) mice lacking (Glu-CB_1_-KO) or not (Glu-CB_1_-WT) CB_1_Rs from cortical glutamatergic neurons using the Nex-Cre recombinase (*12*, *35*, *37*); (vi) mice lacking (GFAP-CB_1_-KO) or not (GFAP-CB_1_-WT) CB_1_Rs from astrocytes (use of the tamoxifen-inducible GFAP-CreErt2 recombinase: (*45*, *46*)), (vii) mice from a knock-in line lacking (DN22-CB_1_-KI) or not (WT) the first 22 amino-acids allowing CB_1_Rs trafficking to mitochondria-associated membranes (*28*, *45*), (viii) mice lacking or not CB_1_Rs from striatal D1-expressing neurons (D1-CB_1_-KO/WT, formerly termed DrD1-CB_1_-KO/WT (*54*), and mice lacking (A2A-CB_1_-KO) or not (A2A-CB_1_-WT) CB_1_Rs from D2-expressing neurons (*54*), and (ix) mice lacking (PV-CB_1_-KO) or not (PV-CB_1_-WT) CB_1_Rs from PV-expressing cells. To respectively re-express CB_1_R in GABAergic neurons and in glutamatergic neurons of mice lacking CB_1_R expression (STOP-CB_1_ mice), GABA-CB_1_-Rescue mice or Glu-CB_1_-Rescue mice were also used (*12*, *41*). The VGAT-CB_1_ line was obtained through crossings between male B6J.129S6(FVB)-Slc32a1^tm2(cre)Lowl^/MwarJ mice (JAX 028862; The Jackson Laboratory, USA) and female CB_1_-floxed mice. The PV-CB_1_ line was obtained through crossings between male B6;129P2-Pvalb^tm1(cre)Arbr^/J mice (JAX 008069; The Jackson Laboratory, USA) and female CB_1_-flox mice. The DAGL-KO line was obtained by crossing female B6J Dppa3^tm1(cre)Peli^ (Stella-Cre) mice established in our breeding facilities and male heterozygous DAGLα-flox mice (*75*) (B6J.Dagla tm1) before successive crossings between heterozygous males and females Stella-DAGLα. All mutant and WT mice from each line were littermates and were regenotyped at the end of the experiments.

### Housing

Mice were given at least a 7-day habituation to a partly reversed 12h/12h light/dark cycle (lights off: 9:00-21:00). During that period, all animals were individually housed (to avoid inter-individual aggression) with food and water *ad libitum* in a thermoregulated room (21-22°C). In one series of experiments, male C57Bl/6N were initially housed in 3 per cage before being placed in 2 per operant chamber during the closed economy protocol (see below). All training and PR sessions occurred during the dark phase of the cycle.

### Operant apparatus

The behavioral set-up comprised 15 individual operant chambers (Imetronic, France) located in a room adjacent to the animal housing room (*12*, *13*). These chambers were connected to a computer which recorded all exercise and feeding variables. For operant running experiments, lateral walls were made of grey Perspex while the rear wall had a central hollow for mounting the 20-cm diameter wheel, the release trigger of which was connected to a circuit enabling the wheel to be locked or unlocked by means of a brake-pad. A cue-light placed above the wheel indicated the wheel unlocking. The wheel was flanked by two small ports with cue lights located above to allow the animal to ‘poke’ its nose through. For operant feeding, the rear side (running wheel, nose poke ports, cue-lights) was covered by grey Perspex whereas the left panel of the chamber housed in its center a recessed pellet tray surrounded by two nose poke ports. Cue-lights were placed above the nose poke ports and the feeder to indicate respectively effectiveness of the nose poke and pellet distribution. Nose poke performance could be either “active” (leading to the illumination of the reward-associated cue-light followed by wheel unlocking or pellet distribution) or “inactive” (having no consequence), the left/right allocation of active/inactive nose poke ports being counterbalanced between animals.

### Operant running

For two daily sessions, mice underwent habituation to the chambers and the pairings between cue light illumination and reward availability (*12*, *13*). Thus, mice were placed in the chambers with the cue light above the unlocked running wheel remaining illuminated while the two nose poke ports were covered-up by tape. When learning sessions began, the wheel locking/unlocking mechanism and the nose poke ports were fully operational. The wheel was unlocked for 60 s following a nose poke executed in its allocated active nose poke port. In the FR1 condition, a single active nose poke was sufficient to simultaneously illuminate the cue-light above the port for 10 s and unlock the running wheel for 60 s under light. Nose pokes in the inactive port were counted but had no consequence. When the 60 s had elapsed, the wheel-light extinguished and the brake applied, so that the mouse had to step down from the wheel and execute a further nose poke in order to unlock it again. Nose pokes in the active port while the wheel was already unlocked were without consequence. Habituation and FR1 sessions were ran once daily and lasted for 60 min. For experiments aimed at testing mice under FR1, FR3 and PR reinforcement schedules, there were 5 FR1 sessions followed by 5 FR3 sessions (wherein a 60-s wheel-running period was contingent on 3 consecutive nose pokes in the active port) (*12*). The day after the last FR3 session, mice were tested under a linear PR schedule of reinforcement where the number of active nose pokes required to free the running wheel was incremented by 3 between each rewarded step, with a time limit of 15 min between two successive steps (*12*). After effective training to the operant running protocol, as assessed by stable performances, mice were then shifted to habituation and training sessions for the operant feeding protocol.

### Operant feeding

Because the closed economy protocol aimed at comparing exercise motivation with motivation for a standard food diet (20-mg pellets; F0163, Bio-Serv, Plexx France), we fasted overnight all mice prior to their initial habituation to the operant feeding protocol. Mice were placed in the chambers with the cue light above the pellet tray remaining illuminated while the 2 nose poke ports were covered-up by tape. Immediately after placement of the mouse in the operant chamber, 17 food pellets were successively distributed to the tray (*12*, *76*). This 30-min session was aimed at habituating the mice to the feeder and the cues indicating pellet distribution. Immediately after this session, the feeder was emptied out (if needed) and the nose poke tapes were removed. Mice were then exposed to a 30-min feeding session (this short duration being selected to avoid satiety: (*12*, *76*)) with the nose poke ports being fully operational. During this session, illumination of the cue-light above the feeder and distribution of one food pellet obeyed FR1 reinforcement schedules with inter-trials of 15-sec duration. As for conditioned running, nose pokes in the inactive port were counted but had no consequence. This session was followed by similar ones (except that mice were again provided *ad libitum* food in their home cages) until stabilization of their daily number of active nose poke visits and significant discrimination between active and inactive nose poke ports. In one series of experiments, sucrose-enriched chocolate-flavored 20-mg pellets (5TUL; Test Diet, Bio-Concept Scientific, France) - instead of standard diet pellets - were used during operant training and the closed economy process.

### Operant exercise/food choice

This step involved the daily placement of the mice in the chambers with the possibility to work for either reward (choice protocol). Thus, animals were placed in a choice condition with either wheel unlocking or food distribution being accessible under an FR1 schedule (*12*). However, choosing one reward excluded the possibility to obtain simultaneously the second reward. To enable enough choice trials during these 60-min training sessions, the duration of activation of the wheel was reduced to 20 s whilst that of the feeder remained 15 s. However, to further indicate to the mice that ran during the entire 20-s sequence that the reward choice was mutually exclusive, we added a 5-s period during which a green ceiling light was switched on whilst none of the nose poke ports was active. In most instances, mice displayed stable active nose pokes for the wheel and the food after 3 choice sessions owing to the abovementioned training efficiency to get either reward when proposed alone under FR1 schedules. The day after the last 1-h choice training session (always performed in *ad libitum* fed mice), mice were placed for 12 days in the operant chamber to be tested under closed economy conditions.

### Permanent closed economy protocol

Mice were weighed in the morning and then housed in the operant chambers with a plastic shelter filled with cotton nesting material and *ad libitum* water access (Fig. 1A). The nesting material was yet provided the day before in the home cage to minimize transfer stress. To ease comfort, the grid floor of the operant cage was replaced by a drawer filled with sawdust. Mice had conditioned accesses to a running wheel (1-min duration) or 3 x 20-mg food pellets in an exclusive manner (the two rewards could not be obtained at the same time) under fixed reinforcement schedules. As opposed to the training sessions, there was no time-out periods between reward gains. Initially, mice obeyed for 3 days FR1 rules before being successively exposed to FR3, FR10, and FR30 schedules of reinforcement (3 days/FR), the protocol thus lasting 12 consecutive days. Note that the nose poke scores reached during the first day of exposure to the closed economy protocol (i.e. first of the 3-FR1 days) were never included in the analyses. Thus, food spillage and/or very high running activity could sometimes be observed during that first day, but not thereafter. Each morning, the mice were removed from the chambers and placed for 15 min in their home cage (without food or water) (i) to measure their body weights and their water consumptions and (ii) to check that all nose poke devices were efficient. If needed, sawdust was added to each drawer. In rare cases, mice were removed from the protocol if they lost 20% of their initial body weights for 2 consecutive days.

### Closed economy variables

Body weight changes were calculated as percent changes from the initial body weights and averaged per FR reinforcement schedule. As for body weight analyses, the daily numbers of nose pokes performed by each mouse, and hence its number of reinforcers, were calculated and then averaged over the 2 (FR1) or 3 (FR3-FR30) consecutive days to provide one mean value/FR schedule. Mean running duration per rewarded sequence (maximum: 1 min) and wheel running preference over feeding (i.e. the ratio of the nose pokes for running over the sum of the respective numbers of nose pokes for running and feeding) were individually calculated and then averaged in a similar manner.

Closed economy settings permit to estimate a key behavioral economic variable, namely the EV for a reward, and thereby to compare this value with that of another reward provided the k parameter is similar for both rewards (*17*, *19*, *20*). This variable is inversely linked to the sensitivity of this reward to the strength of the efforts (i.e. the prices) needed to obtain this reward, and is considered a proxy of the intrinsic value of a reinforcer (*19*). Thus, the harder the individual works for this reward with price increases, the higher is the EV and hence the lower is the so-called “elasticity” of the demand. To calculate this EV, we first log-transformed the mean reinforcer numbers per FR schedule (Q) as to obtain Q_0_ and α values using the formula: log Q = log Q_0_ + k(e-^α.Qo.C^ -1) where Q_0_ is the theoretical number of rewards obtained without any effort prerequisite (FR0), k the constant defining the range of reward numbers (constantly set to 2 throughout the study), C the FR schedule (effort, price), and α the so-called elasticity of the demand (*20*). All EVs, measured using the formula: EV = 1/(100.α.k^1.5^) (*19*, *20*), Q_0_ and P_max_ values (transition points between the elasticity and the inelastivity of the demand for a given reward; (*19*)) were provided on-line (http://hdl.handle.net/1808/14934) by an automated calculator through the courtesy of Kaplan B.A. and Reed D.D. (2014).

### Viral vectors and surgery procedures

Mouse analgesia was achieved with a subcutaneous injection of buprenorphine (Buprecare; 0.05 mg/kg). Then, mice were anaesthetized through 5% isoflurane inhalation and placed into a stereotaxic apparatus (Kopf Instruments Model 900; CA, USA) with mouse adaptor and lateral ear bars, the anesthesia being maintained at 1.5% during the entire surgery. Local analgesia with lidocaine (Lidor; 7 mg/kg) was injected under the skin of the head before incision. AAV vectors were bi-laterally infused through a glass pipette using a microinjector (Nanoject II, Drummond Scientific, PA, USA) at a speed of 3 nl/s. For the local deletion of CB_1_Rs in the VMS (500 nl/side; AP: +1.5; ML: ± 0.75; DV: -4.5) or the VLS (500 nl/side; AP: +0.98; ML: ± 1.80; DV: -4.92), CB_1_-flox mice were locally injected with an AAV-CAG-CRE-GFP (5.2 x 10^10^ vg/ml) or its control AAV-CAG-hrGFP virus (3.0 x 10^10^ vg/ml) (*77*). To remove CB_1_Rs from VMS MSNs, we used the AAV-CaMK-HA-CRE construct (400 nl/side; 1.5 x 10^11^ vg/ml) and its control (AAV-CaMK-hrGFP; 6.7 x 10^10^ vg/ml). To rescue CB_1_Rs in the VMS of GABA-CB_1_-KO mice, we injected (500 nl/side) the AAV-CAG-flex-CB_1_-GFP construct (8.3 x 10^10^ vg/ml) and its control (AAV-CAG-flex; 2.3 x 10^11^ vg/ml) (*77*) using the above mentioned coordinates. All viral constructs were generated by Calcium Phosphate transfection of HEK293T cells and purified as described (*35*). After the surgeries, all mice received i.p. injections of meloxicam (Metacam; 5 mg/kg), before being placed in a ventilated cage maintained at 28°C until full anesthesia recovery. Metacam treatments were again provided for the 3 days which followed surgery. Mice were tested at least 3 weeks after the AAV administration to ensure full recovery (normal body weight, complete healing) and expression of the AAV vector.

### Brain tissue sampling

After the closed economy protocol, mice were deeply anesthetized by subsequent injections of xylazine (Rompun; 20 mg/kg) and pentobarbital (Euthasol; 400 mg/kg) and then transcardially perfused with a phosphate-buffered solution (PBS 0.1M, pH 7.4) followed by 4% paraformaldehyde (PFA; BO501128-4L, Sigma). Brains were extracted and post fixed in 4% PFA overnight at 4°C, and then transferred to a 30% sucrose solution for cryopreservation. Brains were then frozen in isopentane (M32631, Sigma) and stored at -80°C. Serial 30-μm free-floating coronal sections were cut with a cryostat (Leica Biosystems CMI1950S) and stored in an antifreeze solution at -20°C. Some sections were counterstained with 4’,6-diamidino-2-phenylindole (DAPI 1:20000, Fisher Scientific) to visualize cell nuclei and then mounted, dried and cover-slipped.

### Immunohistochemistry (IHC) for light microscopy

#### Immunostaining against PV

Brain slices were washed with PBS and then permeabilized for 1h in a blocking solution (with PBS 1x: 10% goat serum; 0.3% triton X-100) at room temperature (RT). Free-floating sections were incubated with guinea-pig anti-PV (1:1000; 195004, Synaptic systems) and revealed with a secondary goat anti-guinea-pig Alexa 555 antibody (1:500; 4413S, Cell Signaling). Images of these sections were taken by a slide scanner (nanozoomer 2.0 HT, Hamamatsu) and counting of PV-labelled cells was performed manually. Image analysis and counting was performed bilaterally in one 40x images per striatum, restricted to the dorsolateral area, with 8 striatal slices per mouse analyzed.

#### Immunostaining against CB_1_R

CB_1_R immunodetection was performed by incubating slices for 1 h with 10% goat serum and then overnight at 4°C with a rabbit anti-CB_1_ (1:500; AF380, Frontier Institute). After washes with a PBST solution (0.3 % Triton X-100 in PBS 1x), the signal was revealed by incubating 2 h at RT with the secondary goat anti-rabbit antibody (1:500; A11008, Invitrogen). The intrinsic fluorescence of the virus and the re-expression of CB_1_R was visualized with an epifluorescence Leica DM600 microscope and a Leica SP2 microscope 40x objective (Leica, Germany).

#### Immunostaining against HA

Sections were washed with PBST, permeabilized in a blocking solution of 10% goat serum in PBST for 1 h at RT. Then, free-floating sections were incubated overnight at 4°C with a rabbit anti-HA (1:1000; 3724, Cell Signaling). After several washes in PBST, slices were incubated for 2 h with a goat anti-rabbit conjuged to Alexa 594 (1:500; A11012, Invitrogen) and then washed in PBST at RT. The fluorescence was visualized with an epifluorescence Leica DM6000 microscope.

### Double Fluorescent *in-situ* hybridization (FISH)

Double FISH experiments to assess CB_1_ transcripts in NAc D1- and D2-expressing neurons were carried out as previously described (*78*). For the probes, a fluorescein-labeled riboprobe against mouse CB_1_R transcript (CB_1_-FITC; 1:1000) and digoxigenin-labeled riboprobes against mouse D1 or D2 transcripts (D1-DIG or D2-DIG; 1:1000) were used. Free-floating sections were treated with 0.2 M HCl, followed by acetylation with 0.25% acetic anhydride in 0.1M triethanolamine (pH 8) for 10 min. In between all steps, sections were rinsed in PBS containing 0.01% diethylpyrocarbonate (DEPC). CB_1_-FITC and D1-DIG (or D2-DIG) were diluted 1:1000 in hybridization buffer and denaturated at 90°C for 5 min. Sections were hybridized overnight at 57°C with the mixture of probes and subsequently washed at 62°C using different stringency wash buffers. Then, sections were incubated for 1 h in sheep anti-FITC-POD antibody (1:1500; Roche, 11426346910) followed by a tyramide signal amplification (TSA) reaction using Cyanine 5 (Cy5)-labeled tyramide (1:100 for 10 min) (NEL744001KT, Akoya Biosciences). After peroxidase quenching with 3% H_2_O_2_ in PBS (30 min) and treatment with 0.2 M HCl (20 min), sections were incubated overnight at 4°C in anti-DIG-POD antibody (1:1500; 11207733910, Roche). The day after, revelation was made using TSA-Cy3 (1:100 for 10 min) (NEL745001KT Akoya Biosciences).

To assess CB_1_R deletion in GABAergic and PV-positive cells of VGAT-CB_1_-KO mice and PV-CB_1_-KO mice, respectively, double FISH assays were performed using DIG- and FITC-labeled riboprobes (1:1000) against mouse CB_1_R and either GAD65 or PV, respectively. Experiments were conducted according to the protocol described above, except for temperature adjustments in hybridization and post-hybridization. For the VGAT-CB_1_ mice, hybridization and washes were performed at 70°C, whereas for PV-CB_1_ mice, hybridization was performed at 61°C and post-hybridization washes at 67°C. Slices were visualized with either an epifluorescence Leica DM600 microscope or a confocal Leica SP2 microscope. For the quantification of CB_1_-expressing PV cells, counting was performed bilaterally in one (40 x) image per striatum, with 3 striatal slices per mouse analyzed.

### Single FISH combined with IHC

CB_1_R deletion in the VMS or VLS was assessed by immunodetection combined with FISH against CB_1_ transcripts, as previously described (*79*). Briefly, slices were incubated overnight at 4°C with rabbit anti-GFP (1:500) (A11122, Invitrogen). The next day, slices were incubated in goat anti-rabbit HRP (horseradish; 1:500) (7074S, Cell Signaling) for 2 h at RT and a tyramide signal amplification (TSA-biotin; 1:250) reaction was used to increase the signal. After an acetylation step, digoxigenin (DIG)-labeled riboprobes against mouse CB_1_R were prepared (1:1000). After overnight hybridization at 62°C, the slices were washed with different stringency wash buffers at 67°C. The probe was revealed by incubating 1 h at RT the sheep anti-DIG-POD antibody (1:1500; 11207733910, Roche) followed by a tyramide signal amplification reaction using Cyanine 3 (Cy3)-labeled tyramide (1:100 for 10 min) in order to detect CB_1_R. After several washes, sections were incubated 30 min in neutravidin 488 (1:500; 22832, Invitrogen) to reveal GFP signal. Finally, slices were incubated 5 minutes in DAPI before being washed, cover-slipped and visualized with an epifluorescence Leica DM6000 microscope and imaged with a confocal Leica SP2 microscope 40x objective (Leica, Germany). For the control virus experiment (AAV-CAG-hrGFP), a single FISH against CB_1_R was performed using the same probe.

### Quantitative real-time PCR

To quantify CB_1_R transcripts in male/female DAGLα-KO mice, compared to their DAGLα-WT littermates, mice were sacrificed by cervical dislocation and their brains rapidly removed. Brain regions were rapidly dissected out on ice before being rapidly frozen on dry ice. RNA tissue extraction and quantitative real-time PCR were performed as previously reported (*80*). For the determination of the reference genes, the RefFinder method was used. Relative expression analysis was normalized against two reference genes: ATP synthase, H+ transporting mitochondrial F1 complex, beta subunit (*Atp5f1b*) and Elongation factor 1-alpha 1 (*Eef1a1*) were used for the frontal cortex, phosphoglycerate kinase 1 (*Pgk1*) and succinate dehydrogenase complex subunit (*Sdha*) for the striatum, phosphoglycerate kinase 1 (*Pgk1*) and non-POU-domain-containing, octamer binding protein (*Nono*) for the hippocampus, and ATP synthase, H+ transporting mitochondrial F1 complex, beta subunit (*Atp5f1b*) and glyceraldehyde-3-phosphate dehydrogenase (*Gapdh*) for the hypothalamus (for primer sequences, see Table S1).

### Statistics

All statistical analyses were performed using GB-Stat software (version 10; Dynamic Microsystems Inc., Silver Spring), except for Kolmogorov-Smirnov normality tests and Grubb’s test for outliers which were both assessed using GraphPad Prism (version 8; GraphPad, San Diego). All data are shown as mean ± SEM. When normal, data were compared through one-tailed or two-tailed Student’s t-test, one-, two- or three-way analyses of variances (where appropriate) before being followed, only if main factor interactions proved significant, by inter-group comparisons using Tukey tests. When data failed to reach normality, Mann-Whitney U-tests were used for two-group comparisons or Kruskal-Wallis h-tests for multiple comparisons, followed if significant, by *post hoc* Mann-Whitney U-tests for inter-group comparisons. In some instances, groups with heterogeneities of the variances were compared following a log-transformation.

## Acknowledgments

The authors thank all the personnel of the Animal Facilities, of the Genotyping platform, and of the Transcriptome platform of the Neurocentre Magendie INSERM U1215. We also thank Dr Vincent Simon (team Energy Balance and Obesity led by Dr Daniel Cota, INSERM U1215) for his useful advices for some fluorescent *in situ* hybridization assays, and Dr L. Di Lodovico (CMME, GHU Paris Psychiatrie, Paris, France) for help during the behavioral experiments. The authors acknowledge the help of Dr A. Zimmer (University of Mainz, Mainz, Germany) who provided us with the DAGLα-flox mice(*75*) (B6J.Dagla tm1). Part of the microscopy work was performed in the Bordeaux Imaging Center, a service unit of the CNRS-INSERM and Bordeaux University, member of the national infrastructure France BioImaging supported by the French National Research Agency (ANR-10-INBS-04).

## Funding

INSERM (GM)

European Research Council (Micabra, ERC-2017-AdG-786467; GM)

Fondation pour la Recherche Médicale (DRM20101220445; GM),

Région Aquitaine (CanBrain, AAP2022A-2021-16763610 and -17219710; GM)

University of Bordeaux IdEx GPR Brain 2030 (Neurocentre Magendie platforms)

IReSP (AAP-IReSP-Addiction 2023; FC).

## Author contributions

Study design: IH, RF, BR, GM, FC

*In vivo* and *ex vivo* experiments: IH, RF, BR, AEP, FC

Histochemistry and fluorescent *in situ* hybridization: IH, RF, DG, FJ-K

Plasmids and AAVs: AC, LB

Quantitative real-time PCRs: TL.-L

Draft writing: FC

Review and editing: all authors

## Data and materials availability

All data needed to evaluate and reproduce the results are present in the paper and/or the Supplementary Materials.

