## Supplementary Figures S1 to S9, Table S1 for "Ventromedial striatal GABAergic interneurons sex-dependently gate cost-benefit choices between food and exercise"

**This PDF file includes:**

Figs. S1 to S9

Table S1

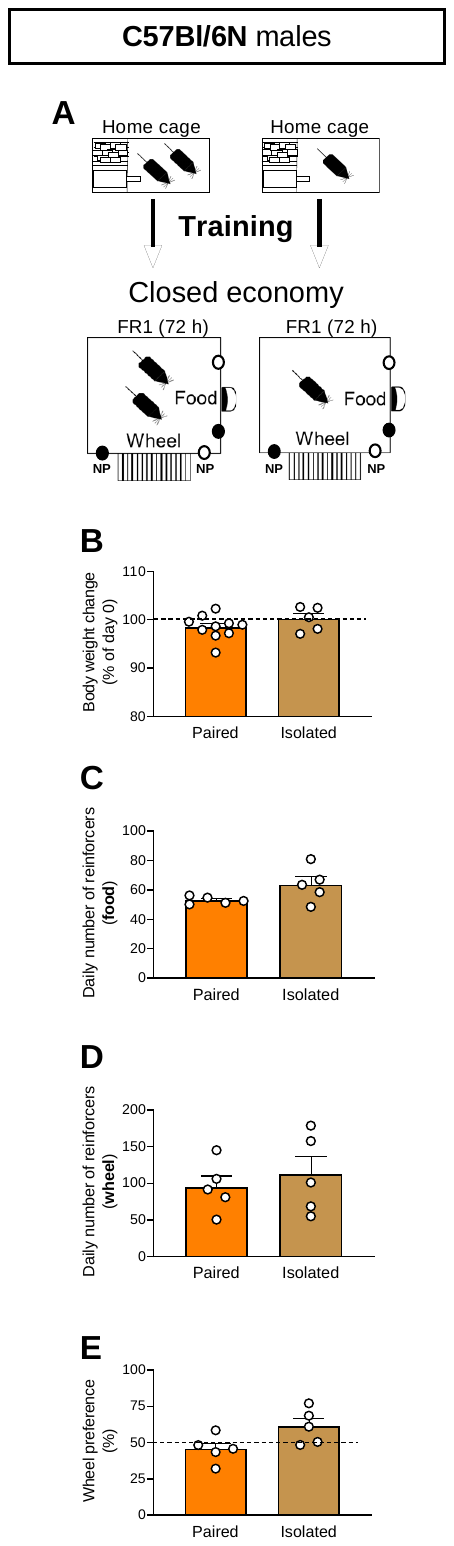

**Fig. S1.** (**A**) Scheme of the procedure with pair- and single-housed C57Bl6/N male mice. (**B**) Pair-housed (n = 5 pairs) and single-housed (n = 10) mice do not differ in their body weight changes during the 3-day exposure to the closed economy protocol (two-tailed Student t-test p = 1.25, NS). (**C, D**) Paired- and single-housed mice do not differ respectively in their daily numbers of food and wheel reinforcers (**C**, two-tailed Student t-test p = 1.96, NS; **D**, two-tailed Student t-test p = 0.60, NS). (**E**) The wheel preference over feeding is similar in pair- and single-housed mice (two-tailed Student t-test p = 2.25, NS). Except for individual body weight measures, all other variables were calculated as means per chamber (n = 5). All data represent mean ± SEM.

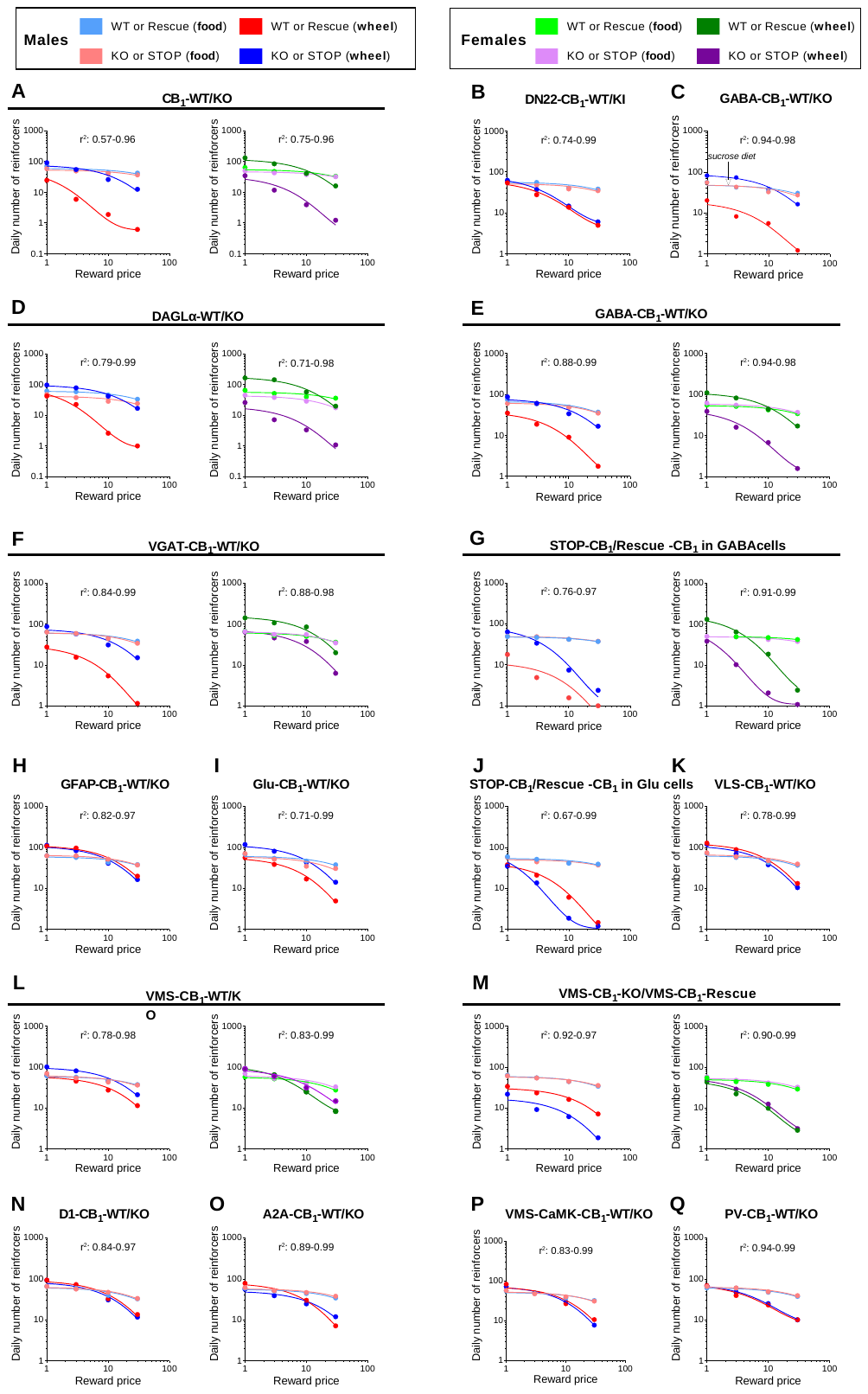

**Fig. S2.** (**A-Q**) Food and exercise demand elasticity curves with their best fit R squared values for all the experiments reported in this study. For the first series of experiments using C57BL/6N mice, demand curves are shown under Fig. 1I.

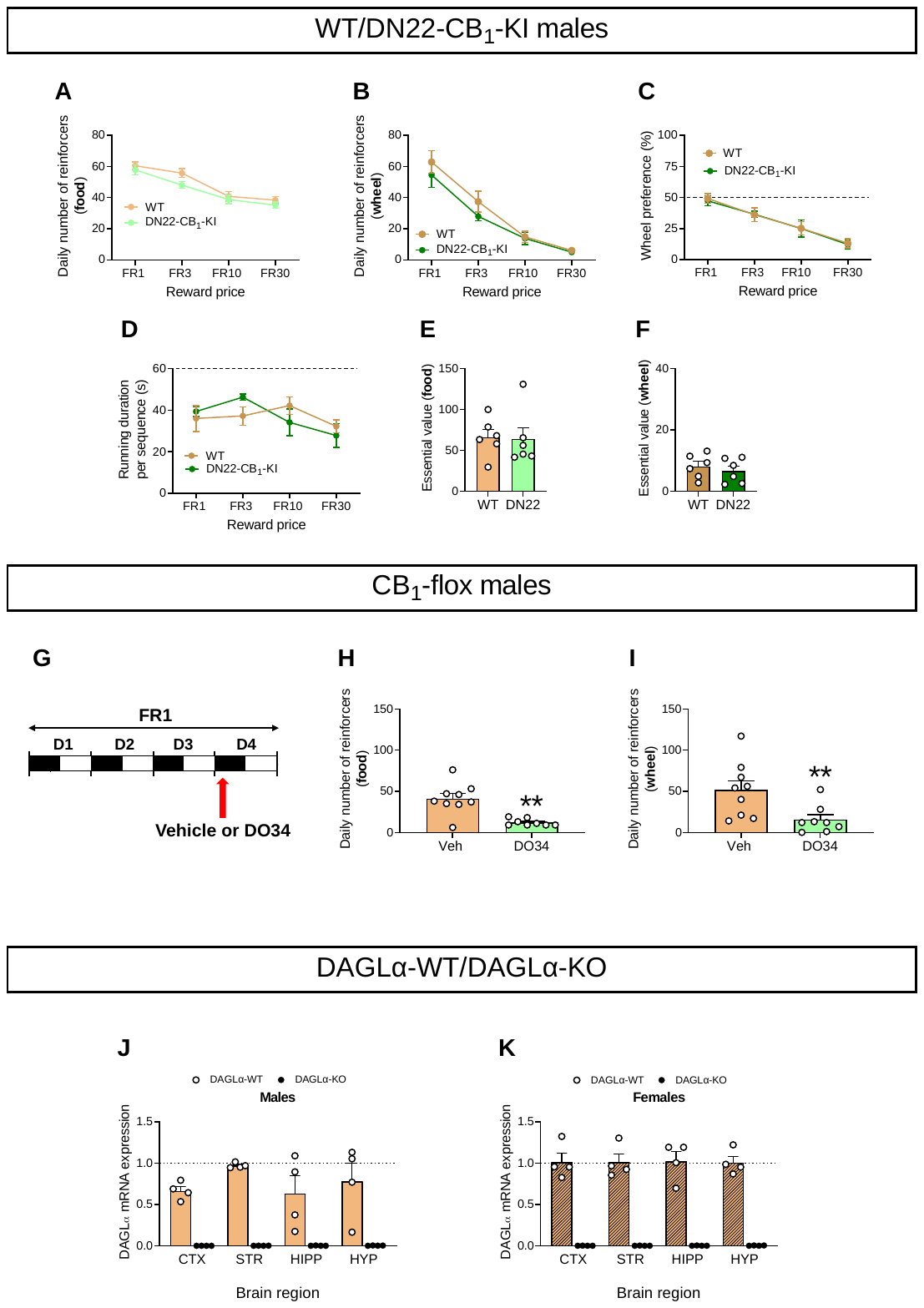

**Fig. S3.** (**A, B**) Daily numbers of food and exercise reinforcers achieved by male WT and DN22-CB_1_-KI mice (n = 6/genotype) for each FR schedule (**A**, two-way repeated ANOVA genotype effects: p = 0.12, NS; **B** two-way repeated ANOVA genotype effects: p = 0.18, NS). (**C**) Wheel preference over feeding (two-way repeated ANOVA genotype effects: p = 0.87, NS). (**D**) Running duration (maximum: 60 sec) for each rewarded sequence of wheel access (two-way repeated ANOVA genotype effects: p = 0.99, NS). (**E, F**) Food and exercise essential values in WT and DN22-CB_1_-KI mice (**E**, two-tailed Student t-test p = 0.89, NS; **F**, two-tailed Student t-test p = 0.56, NS). (**G**) Protocol of DO34 injection (white and dark zones refer respectively to the light and dark phases of the circadian cycle). (**H, I**) Daily numbers of food and exercise reinforcers achieved by vehicle-injected (n = 9) and DO34-injected (n = 8) CB_1_-floxed mice under FR1 schedules of reinforcement (**H**, Mann-Whitney test: p = 0.0067; **I**, Mann-Whitney test: p = 0.0052). (**J, K**) DAGLα mRNA expression in the frontal cortex (CTX), striatum (STR), hippocampus (HIPP), and hypothalamus (HYP) of male and female DAGLα-WT and DAGLα-KO mice (n = 4/genotype). All data represent mean ± SEM; **p < 0.01.

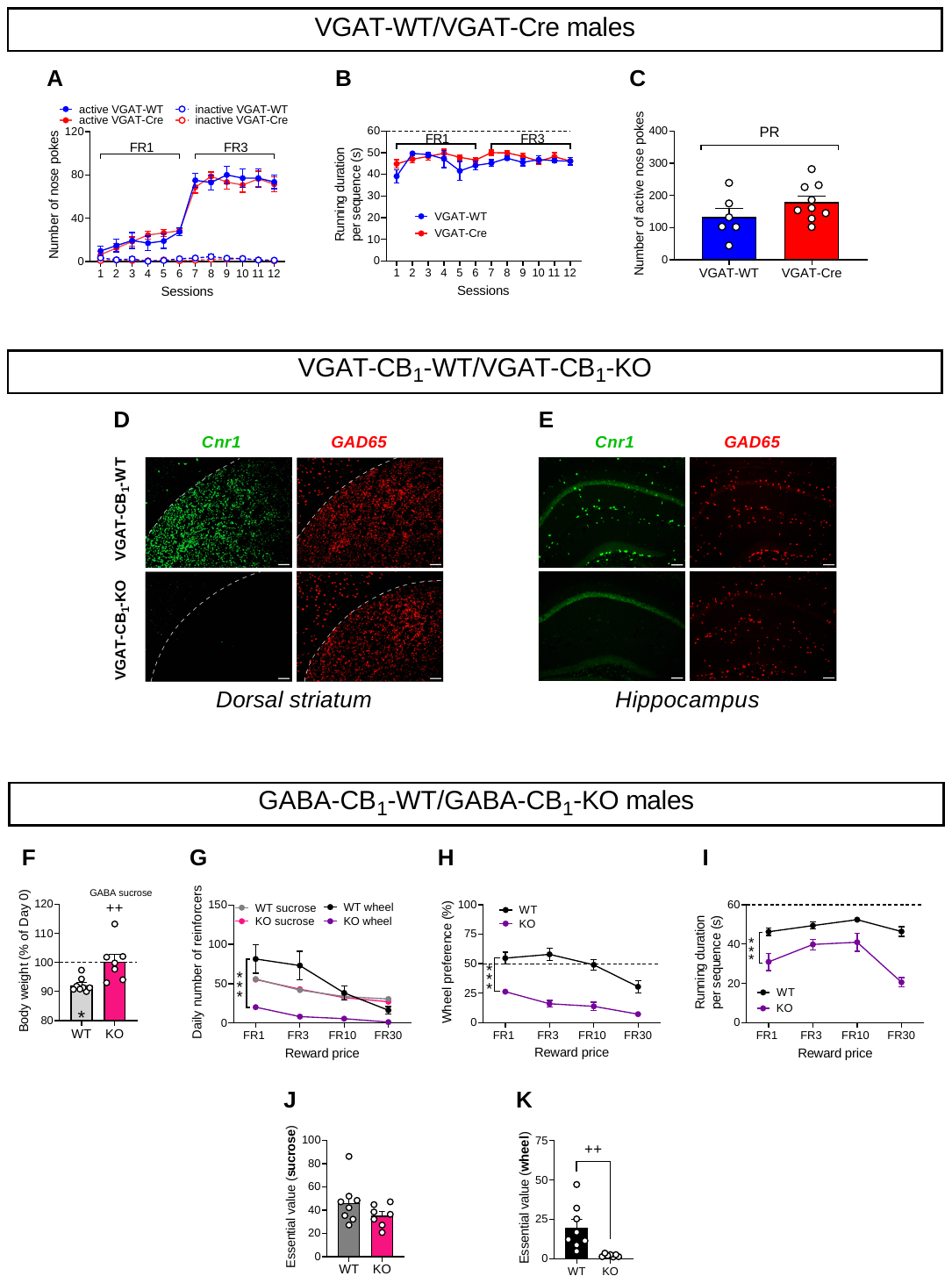

**Fig. S4.** (**A, B**) Nose poke performances and running durations per sequence under FR1 and FR3 reinforcement schedules displayed by male VGAT-WT (n = 6) and VGAT-Cre (n = 9) mice during 60-min sessions (**A**, two-way repeated ANOVA genotype effects: p = 0.91, NS; **B**, two-way repeated ANOVA genotype effects: p = 0.18, NS). (**C**) Number of active nose pokes performed by VGAT-WT and VGAT-Cre males during a 60-min PR session (two-tailed Student t-test p = 0.17, NS). (**D, E**) Representative double fluorescence *in situ* hybridization assays of *Cnr1* and *GAD65* transcripts in the dorsal striatum and hippocampus of male VGAT-CB_1_-WT and VGAT-CB_1_-KO mice (scale bar = 50 µm). (**F**) Percent changes in body weight during the close economy protocol in male GABA-CB_1_-WT (n = 8) and GABA-CB_1_-KO (n = 7) mice offered the choice between a sucrose isocaloric diet and exercise (two-tailed Student t-test: p = 0.0075; one-tailed Student t-test against 100: p < 0.0001 and p = 0.93, NS for male GABA-CB_1_-WT and GABA-CB_1_-KO mice, respectively). (**G**) Daily numbers of food (sucrose diet) and exercise reinforcers achieved by male GABA-CB_1_-WT and GABA-CB_1_-KO mice (two-way repeated ANOVA genotype effects: p = 0.73, NS and p < 0.0001 for food and exercise, respectively). (**H**) Wheel preference over sucrose feeding in male GABA-CB_1_-WT and GABA-CB_1_-KO mice (two-way repeated ANOVA genotype effects: p < 0.0001). (**I**) Running duration (maximum: 60 sec) for each rewarded sequence of wheel access by male GABA-CB_1_-WT and GABA-CB_1_-KO mice (two-way repeated ANOVA genotype effects: p = 0.0001). (**J, K**) Food and exercise essential values in male GABA-CB_1_-WT and GABA-CB_1_-KO mice (two-tailed Student t-test: p = 0.91, NS and p = 0.006, respectively). All data represent mean ± SEM; *p < 0.05, **p < 0.01, ***p < 0.001 and ^++^p < 0.01.

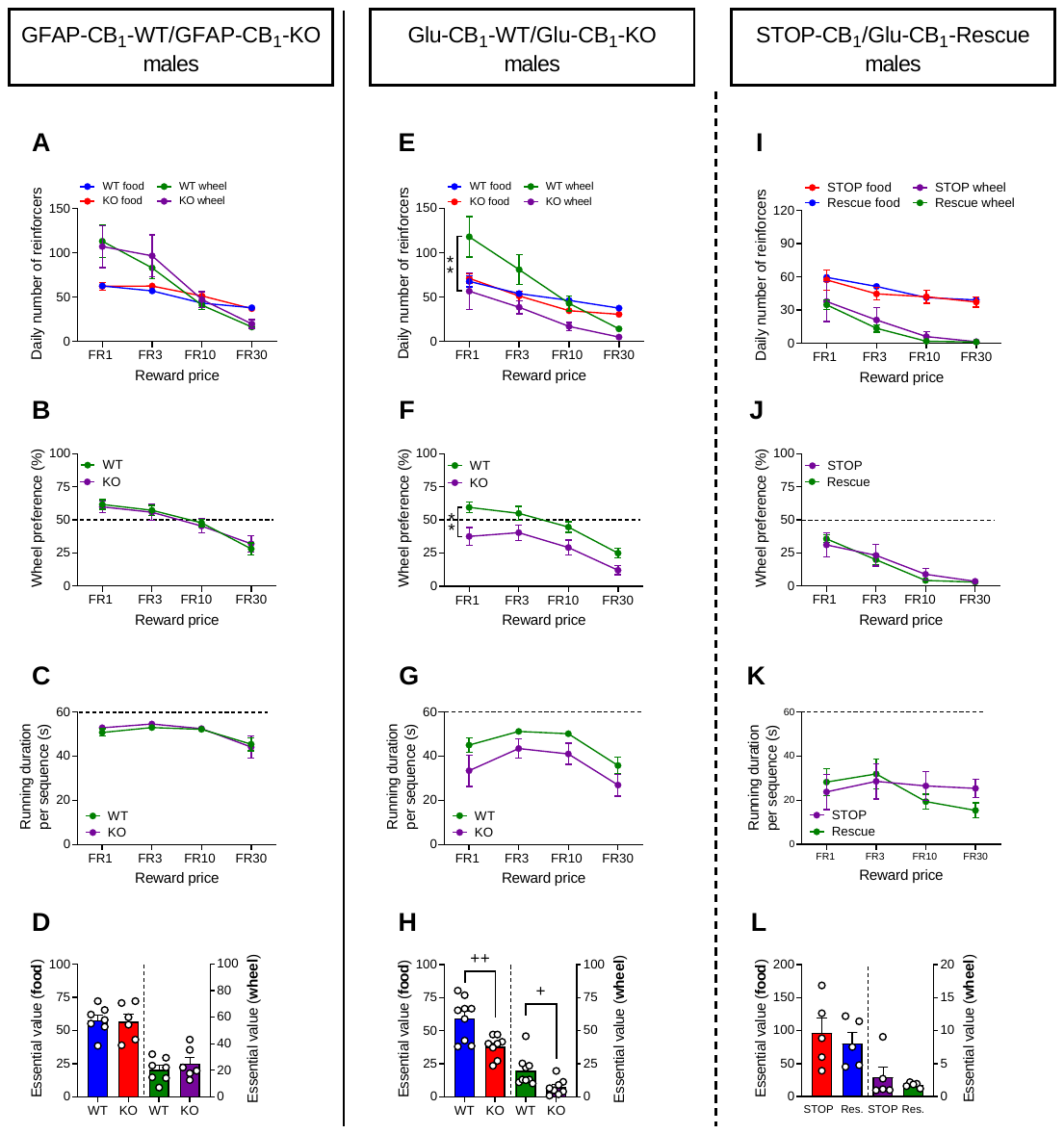

**Fig. S5.** (**A**) Daily numbers of food and exercise reinforcers achieved by male GFAP-CB_1_-WT (n = 7) and GFAP-CB_1_-KO mice (n = 6) for each FR schedule (two-way repeated ANOVA genotype effects: p = 0.28 and p = 0.79 respectively, NS; (**B**) Wheel preference over feeding in GFAP-CB_1_-WT and GFAP-CB_1_-KO mice (two-way repeated ANOVA genotype effects: p = 0.93, NS). (**C**) Running duration (maximum: 60 sec) for each rewarded sequence of wheel access by GFAP-CB_1_-WT and GFAP-CB_1_-KO mice (two-way repeated ANOVA genotype effects: p = 0.71, NS). (**D**) Food and exercise essential values in GFAP-CB_1_-WT and GFAP-CB_1_-KO mice (two-tailed Student t-test p = 0.86 and p = 0.44 respectively, NS). (**E**) Daily numbers of food and exercise reinforcers achieved by male GLU-CB_1_-WT (n = 9) and GLU-CB_1_-KO mice (n = 8) for each FR schedule (two-way repeated ANOVA genotype effects: p = 0.25, NS and p = 0.008 respectively). (**F**) Wheel preference over feeding in GLU-CB_1_-WT and GLU-CB_1_-KO mice (two-way repeated ANOVA genotype effects: p = 0.0097). (**G**) Running duration (maximum: 60 sec) for each rewarded sequence of wheel access by GLU-CB_1_-WT and GLU-CB_1_-KO mice (two-way repeated ANOVA genotype effects: p = 0.07, NS). (**H**) Food and exercise essential values in GLU-CB_1_-WT and GLU-CB_1_-KO mice (two-tailed Student t-test p = 0.0044 and p = 0.016 respectively). (**I**) Daily numbers of food and exercise reinforcers achieved by male STOP-CB_1_ and GLU-CB_1_-Rescue mice (n = 5/genotype) for each FR schedule (two-way repeated ANOVA genotype effects: p = 0.66 and p = 0.69 respectively, NS). (**J**) Wheel preference over feeding in STOP-CB_1_ and GLU-CB_1_-Rescue mice (two-way repeated ANOVA genotype effects: p = 0.86, NS). (**K**) Running duration (maximum: 60 sec) for each rewarded sequence of wheel access by STOP-CB_1_ and GLU-CB_1_-Rescue mice (two-way repeated ANOVA genotype effects: p = 0.76, NS). (**L**) Food and exercise essential values in STOP-CB_1_ and GLU-CB_1_-Rescue mice (two-tailed Student t-test p = 0.50 and p = 0.48 respectively, NS). All data represent mean ± SEM; **p < 0.01 and ^+^p < 0.05, ^++^p < 0.01.

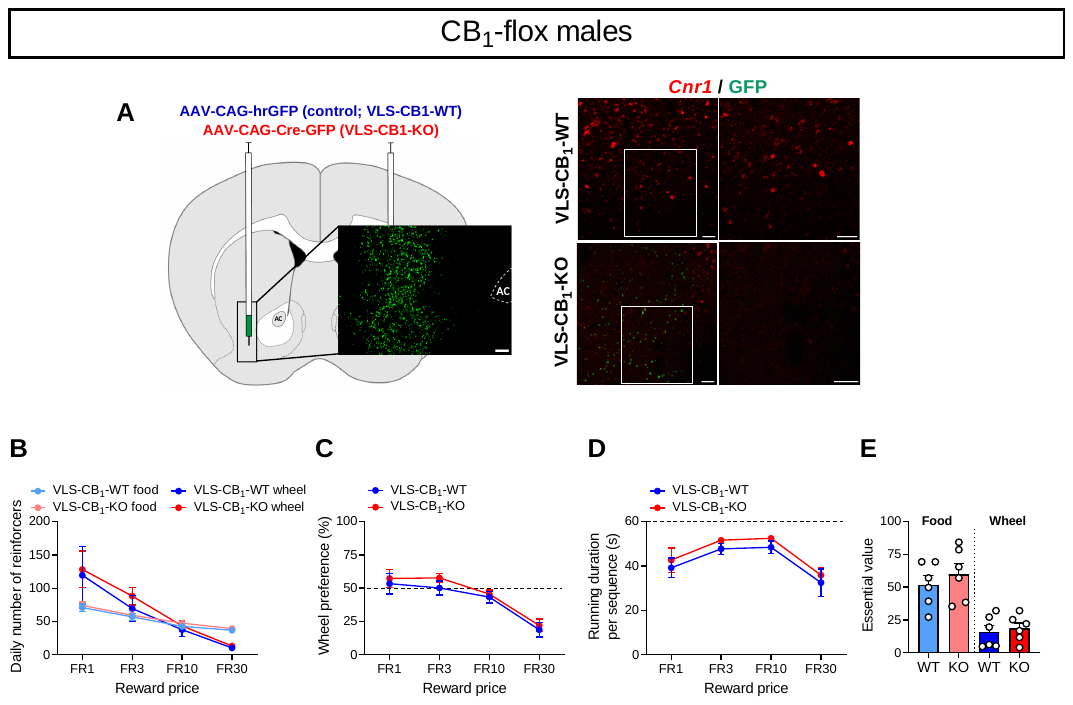

**Fig. S6.** (**A**) Viral-based strategy to respectively delete CB_1_ receptors from the ventrolateral (VLS) part of the striatum of male CB_1_-floxed mice (scale bar = 100 µm) with fluorescence *in situ* hybridization for *Cnr1* transcripts and GFP immunostainings (scale bars = 50 µm). (**B**) Daily numbers of food and exercise reinforcers achieved by male mice injected with the inactive virus (VLS-CB_1_-WT, n = 6) and the active virus (VLS-CB_1_-KO, n = 6) for each FR schedule (two-way repeated ANOVA virus effects: p = 0.50 and p = 0.69 for food and exercise, respectively; NS). (**C**) Wheel preference over feeding in VLS-CB_1_-WT and VLS-CB_1_-KO mice (two-way repeated ANOVA virus effects: p = 0.54, NS). (**D**) Running duration (maximum: 60 sec) for each rewarded sequence of wheel access by VLS-CB_1_-WT and VLS-CB_1_-KO mice (two-way repeated ANOVA virus effects: p = 0.41, NS). (**E**) Food and exercise essential values in VLS-CB_1_-WT and VLS-CB_1_-KO mice (two-tailed Student t-test p = 0.41 and p = 0.67 respectively, NS). All data represent mean ± SEM.

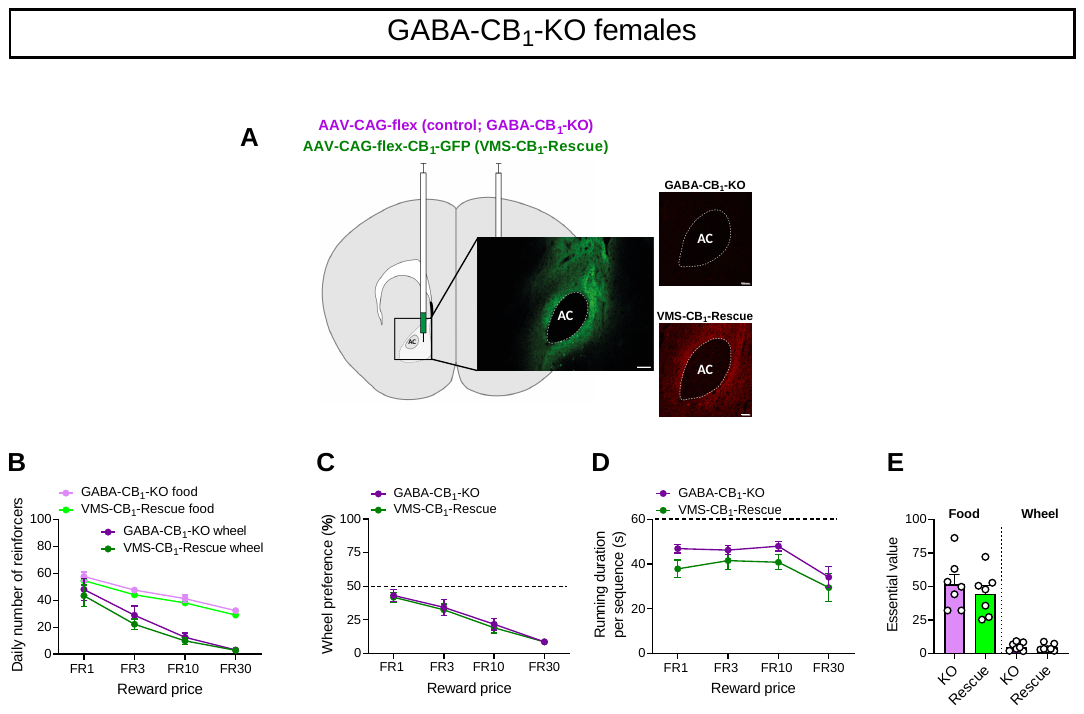

**Fig. S7.** (**A**) Viral-based strategy to re-express CB1 receptors in the VMS of GABA-CB_1_-KO female mice with representative images of GFP immunostainings (scale bar = 100 µm) and CB_1_R transcripts (scale bars = 50 µm) in female mice injected with either the control virus or the rescue virus. (**B**) Daily numbers of food and exercise reinforcers achieved by female mice injected with the inactive virus (GABA-CB_1_-KO, n = 7) and the active virus (VMS-CB_1_-Rescue, n = 7) for each FR schedule (two-way repeated ANOVA virus effects: p = 0.14 and p = 0.50 for food and exercise, respectively; NS). (**C**) Wheel preference over feeding in female GABA-CB_1_-KO and VMS-CB_1_-Rescue mice (two-way repeated ANOVA virus effects: p = 0.74, NS). (**D**) Running duration (maximum: 60 sec) for each rewarded sequence of wheel access by female GABA-CB_1_-KO and VMS-CB_1_-Rescue mice (two-way repeated ANOVA virus effects: p = 0.088, NS). (**E**) Food and exercise essential values in female GABA-CB_1_-KO and VMS-CB_1_-Rescue mice (food: two-tailed Student t-test p = 0.74, NS; exercise: Mann-Whitney test: p = 0.70, NS). All data represent mean ± SEM.

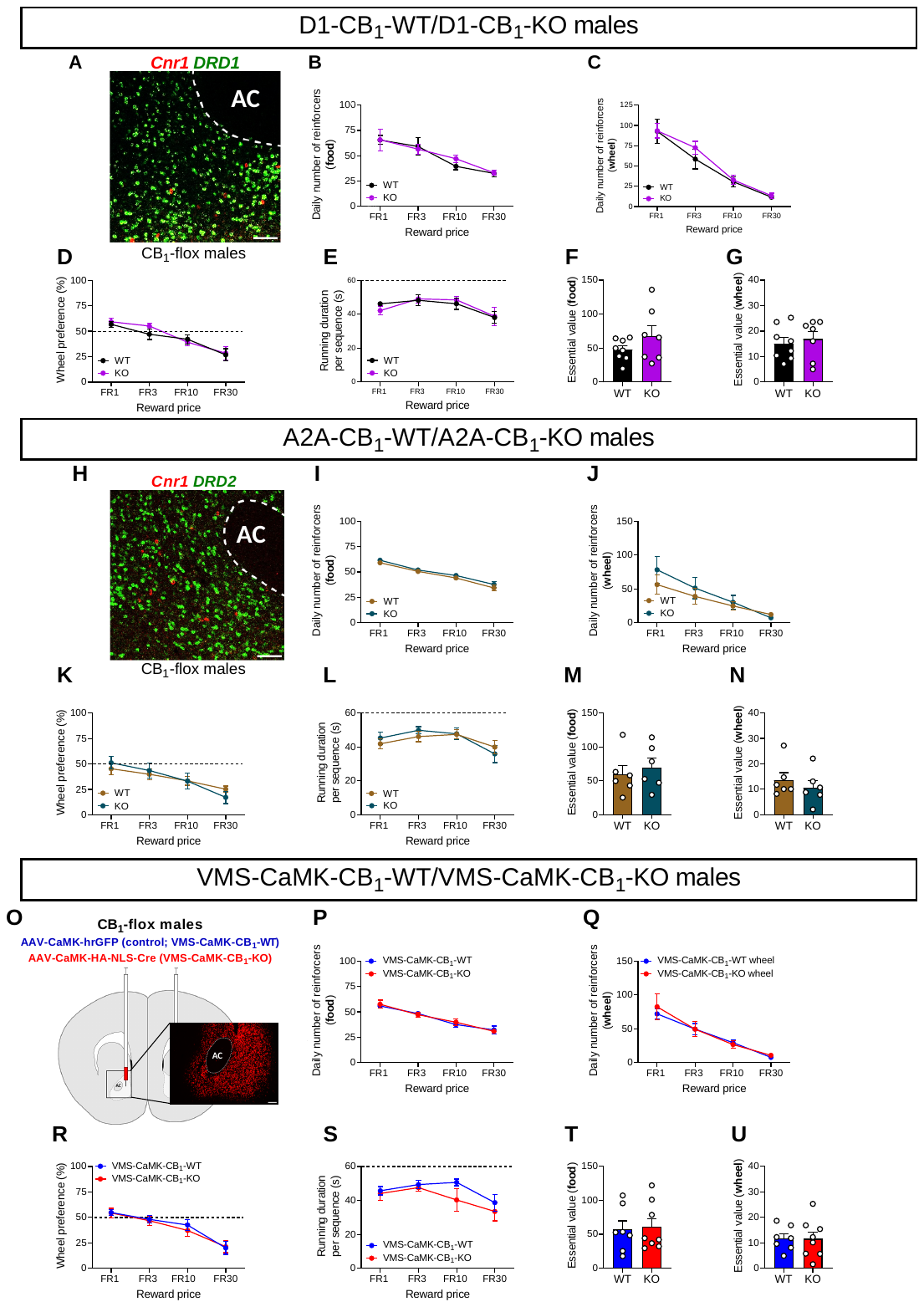

**Fig. S8.** (**A**) Representative fluorescence *in situ* hybridization for *Cnr1* and *DRD1* transcripts in the NAc of one CB_1_-flox male (scale bar = 50 µm). (**B, C**) Daily numbers of food and exercise reinforcers achieved by male D1-CB_1_-WT (n = 8) and D1-CB_1_-KO mice (n = 7) for each FR schedule (**B**, two-way repeated ANOVA genotype effects: p = 0.79, NS; **C** two-way repeated ANOVA genotype effect: p = 0.64, NS). (**D**) Wheel preference over feeding in D1-CB_1_-WT and D1-CB_1_-KO mice (two-way repeated ANOVA genotype effects: p = 0.56, NS). (**E**) Running duration (maximum: 60 sec) for each rewarded sequence of wheel access by D1-CB_1_-WT and D1-CB_1_-KO mice (two-way repeated ANOVA genotype effects: p = 0.99, NS). (**F, G**) Food and exercise essential values in D1-CB1-WT and D1-CB_1_-KO mice (two-tailed Student t-tests p = 0.21 and p = 0.68, NS respectively). (**H**) Representative fluorescence *in situ* hybridization for *Cnr1* and *DRD2* transcripts in the NAc of one CB_1_-flox male (scale bar = 50 µm). (**I, J**) Daily numbers of food and exercise reinforcers achieved by male A2A-CB_1_-WT and A2A-CB_1_-KO mice (n = 6/genotype) for each FR schedule (**I**, two-way repeated ANOVA genotype effects: p = 0.31, NS; **J**, two-way repeated ANOVA genotype effect: p = 0.55, NS). (**K**) Wheel preference over feeding in A2A-CB_1_-WT and A2A-CB_1_-KO mice (two-way repeated ANOVA genotype effects: p = 0.97, NS). (**L**) Running duration (maximum: 60 sec) for each rewarded sequence of wheel access by A2A-CB_1_-WT and A2A-CB_1_-KO mice (two-way repeated ANOVA genotype effects: p = 0.52, NS). (**M, N**) Food and exercise essential values in A2A-CB_1_-WT and A2A-CB_1_-KO mice (two-tailed Student t-tests p = 0.58 and p = 0.74, NS respectively). (**O**) Viral-based strategy to delete CB_1_Rs from ventromedial (VMS) principal neurons (i.e. MSNs) of male CB1-floxed mice with a representative HA immunostaining (scale bar = 100 µm). (**P, Q**) Daily numbers of food and exercise reinforcers achieved by male mice injected with the inactive virus (VMS-CaMK-CB_1_-WT, n = 7) and the active virus (VMS-CaMK-CB_1_-KO, n = 8) for each FR schedule (two-way repeated ANOVA virus effects: p = 0.90 and, p = 0.80, NS respectively). (**R**) Wheel preference over feeding in VMS-CaMK-CB_1_-WT and VMS-CaMK-CB_1_-KO mice (two-way repeated ANOVA virus effects: p = 0.77, NS). (**S**) Running duration (maximum: 60 sec) for each rewarded sequence of wheel access by VMS-CaMK-CB_1_-WT and VMS-CaMK-CB_1_-KO mice (two-way repeated ANOVA virus effects: p = 0.69, NS). (**T, U**) Food and exercise essential values in VMS-CaMK-CB_1_-WT and VMS-CaMK-CB_1_-KO mice (two-tailed Student t-tests p = 0.85, and p = 0.97, NS respectively). All data represent mean ± SEM.

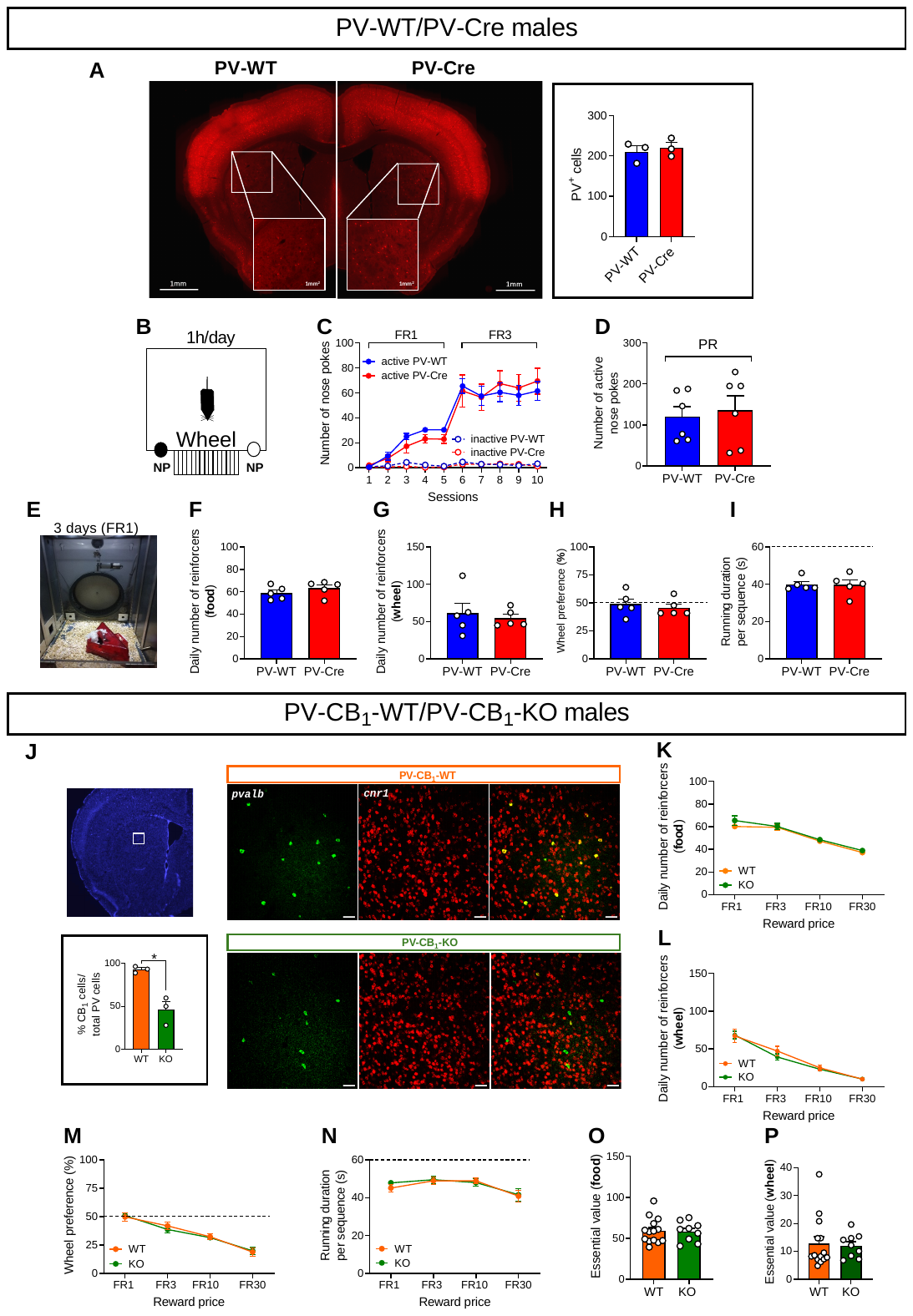

**Fig. S9.** (**A**) Representative immunostainings (with enlarged insets) and numbers (n = 3 mice/genotype; Mann-Whitney test p = 0.99, NS) of parvalbumin (PV)-expressing cells in the dorsal striatum of male PV-WT and PV-Cre mice. (**B**) Protocol to measure the numbers of wheel reinforcers under FR1, FR3 and PR schedules of reinforcements in PV-WT and PV-Cre mice (n = 6/genotype). (**C**) Respective numbers of active/inactive nose pokes under FR1 and FR3 schedules in PV-WT and PV-Cre mice (two-way repeated ANOVA genotype effects p = 0.92, NS). (**D**) Active nose pokes performed by PV-WT and PV-Cre mice during a PR session (two-tailed Student t-test p = 0.74, NS). (**E**) Closed economy protocol to measure food and wheel demands under FR1 reinforcement schedules in PV-WT and PV-Cre mice (n = 6/genotype). (**F, G**) Daily numbers of food and exercise reinforcers achieved by male PV-WT and PV-Cre mice (two-tailed Student t-test p = 0.36 and p = 0.71, respectively; NS). (**H**) Wheel preference over feeding in male PV-WT and PV-Cre mice (two-tailed Student t-test p = 0.74, NS). (**I**) Running duration (maximum: 60 sec) for each rewarded sequence of wheel access by male PV-WT and PV-Cre mice (two-tailed Student t-test p = 0.95, NS). (**J**) Fluorescence *in situ* hybridization for *pvalb* and *cnr1* transcripts (scale bars = 50 µm) and quantification of the respective percentages of CB1-expressing PV cells in the dorsal striatum of male PV-CB_1_-WT and PV-CB_1_-KO mice (n = 3/genotype; Mann-Whitney test p = 0.0495). (**K, L**) Daily numbers of food and exercise reinforcers achieved by male PV-CB_1_-WT (n = 14) and PV-CB_1_-KO (n = 9) mice for each FR schedule (**K**, two-way repeated ANOVA genotype effects: p = 0.23; **L**, two-way repeated ANOVA genotype effects: p = 0.73, NS). (**M**) Wheel preference over feeding in PV-CB_1_-WT and PV-CB_1_-KO mice (two-way repeated ANOVA genotype effects: p = 0.90, NS). (**N**) Running duration (maximum: 60 sec) for each rewarded sequence of wheel access by PV-CB_1_-WT and PV-CB_1_-KO mice (two-way repeated ANOVA genotype effects: p = 0.76, NS). (**O, P**) Food and exercise essential values in PV-CB_1_-WT and PV-CB_1_-KO mice (two-tailed Student t-tests p = 0.97 and p = 0.79, NS respectively). All data represent mean ± SEM.

| **Gene** | **GenBank ID** | **Forward Sequence (5′-3′)** | **Reverse Sequence (5′-3′)** |
| --- | --- | --- | --- |
| *Dagla* | NM_198114 | GGTCCTGCTCGTGCTGTCTC | TGCAGCCACAACAGTTGTCTTC |
| *Atp5f1b* | NM_016774 | GCCCGAGGAGTGCAGAAAA | TCCATACCCAAGATGGCAATG |
| *Eef1a1* | NM_010106 | TGAACCATCCAGGCCAAATC | GCATGCTATGTGGGCTGTGT |
| *Pgk1* | NM_008828 | CCACAGAAGGCTGGTGGATT | TCTGGACTCTCCAAAGCCTTG |
| *Sdha* | NM_023281 | TACAAAGTGCGGGTCGATGA | TGTTCCCCAAACGGCTTCT |
| *Nono* | NM_023144 | CTGTCTGGTGCATTCCTGAACTAT | AGCTCTGAGTTCATTTTCCCATG |
| *Gapdh* | NM_008084 | TCAAGAAGGTGGTGAAGCAG | TGGGAGTTGCTGTTGAAGTC |

**Table S1.** Primer sequences used for the determination of the *Dagla* gene and the reference genes.
